# Male Age and Sexual Maturity: Lipopolysaccharide-induced tumor necrosis factor influences sperm quality and reproduction in *Anopheles culicifacies*

**DOI:** 10.64898/2026.08.31.748190

**Authors:** Pooja Rohilla, Vaishali Saini, Vartika Srivastava, Pooja Yadav, Nirmala Sankhala, Tanvi Singh, Gunjan Sharma, Gitanjali Tandon, Suchi Tyagi, Jyoti Rani, Rajnikant Dixit

**Affiliations:** Laboratory of Host-Parasite Interaction Studies, Department of Vector Genomics, ICMR-National Institute of Malaria Research, Dwarka, New Delhi, India; Academy of Scientific and Innovative Research (AcSIR), Ghaziabad, Uttar Pradesh, India

**Keywords:** Anopheles culicifacies, Male Fertility, Sperm clearance, Testes homeostasis, Phagocytosis, LITAF, Apoptotic Regulation, Reproductive physiology

## Abstract

Elucidating the biological and molecular mechanisms that govern male fertility and mating behavior in mosquitoes is critical for optimizing genetic and sterile insect technique-based vector control strategies. Here, we examined age-related changes in male reproductive capacity in *Anopheles culicifacies*, using female egg output as an indirect indicator of male fertility. Our results demonstrated that male reproductive age follows a non-linear pattern of fertility. Morphometric analysis from emergence to day 13 post-eclosion revealed a strong correlation between seminal vesicle capacity and female fecundity, suggesting that age-dependent gonadal development directly influences reproductive potential. At the molecular level, we identified *AcLITAF6* as a key regulator of male reproductive homeostasis. RNAi-mediated knockdown of *AcLITAF6* impaired apoptosis-associated and phagocytic clearance, reduced sperm viability, and decreased female productive outcomes.

Conclusively, we reveal a previously unrecognized role of LITAF in sperm quality control and male reproductive fitness, highlighting *AcLITAF6* as a potential target for mosquito population suppression strategies.

## Introduction

Mosquito reproductive success is traditionally studied from the perspective of females, as they play a direct role in disease transmission. Yet, male fertility and its regulation are equally pivotal for sustaining vector populations and ensuring the success of emerging control technologies such as the Sterile Insect Technique (SIT) and gene drives [^1–3^]. Decisions on the number and age of modified males to be released are often speculative because little is known about the age-dependent dynamics of male reproductive fitness in natural populations. While post-mating responses in females are well characterized, the physiological and behavioral maturation of males across their reproductive lifespan remains poorly understood [^4,5^].

Sexual maturation in male mosquitoes involves both anatomical and physiological transformations. These primarily include genitalia rotation, testes enlargement, and complete spermatogenesis, while accessory glands begin producing seminal fluids that are critical for sperm function and post-mating responses in females. In *Anopheles gambiae* and *Aedes aegypti*, this process typically requires 24–48 hours post-eclosion under laboratory conditions [^6,7^]. However, environmental factors such as temperature, nutrition, and density can modulate this timeline in the field [^8^]. Importantly, reproductive outcomes are shaped by age: young males may be physiologically immature and less competitive in swarms [^9^], whereas older males often experience declining sperm quality and reduced energy for mating [^10,11^]. These age-related changes create a dynamic landscape of male reproductive potential, where peak competitiveness is tightly linked to the synchrony of physiological and behavioral traits.

The quick-to-court (qtc) gene, first described in *Drosophila*, influences sex-specific behaviors in mosquitoes. In *An. culicifacies*, it shows sexually dimorphic expression, enriched in male olfactory and reproductive tissues. Its activity links male courtship drive to physiological readiness, optimizing mating behavior [^12^]. Thus, reproductive success depends on synchrony between gonadal maturation, sperm replenishment, and mating activity, and mismatches—either between partners or between somatic and germline aging—can impair fertility [^13,14^].

At the cellular level, age-associated reproductive decline is often accompanied by the accumulation of defective or apoptotic germ cells, which must be efficiently removed to preserve testicular homeostasis. This process relies on a coordinated interplay between apoptotic signaling and phagocytic clearance. In *Drosophila*, cyst cells eliminate defective germ cells *via* phagoptosis [^15^], and in *An. gambiae*, the complement-like protein TEP1 eliminates defective spermatogonia, highlighting the integration of immune surveillance with germline quality control [^16^]. Testis-specific expression of antioxidant and immune genes is also known to influence spermatogenesis and sperm longevity [^17^]. Understanding the functional correlation of genes controlling testicular physiological dynamics and sperm quality is crucial for male-mediated female sterility; yet, such studies are currently limited to the model species, such as *An. gambiae* and *An. stephensi* [^18,19^]

While *Anopheles stephensi* and *Anopheles culicifacies* are the dominant malaria vectors in India, account for approximately 20% and 65% of malaria transmission in urban and rural settings, respectively. These species exhibit significant morphological and biological differences in behavioral adaptation, insecticide resistance, and vector competence; however, the molecular basis of these variations remains poorly characterized. A distinguishing feature of *An. culicifacies* is its existence as a complex of five sibling species (A–E), which exhibit marked differences in fecundity, mating behavior, longevity, insecticide resistance, and vector competence despite being morphologically indistinguishable [^20^]. This reproductive and ecological heterogeneity is rarely observed to the same extent in other major malaria vectors and likely contributes to differences in malaria transmission efficiency across endemic regions. These unique characteristics make *An. culicifacies* important for investigating the molecular pathways that regulate reproductive fitness and influence vectorial success.

Recent advances in functional genomics have begun to bridge these knowledge gaps by exploring the unique molecular architecture of adult female *An. culicifacies* [^21–25^]. However, despite recognizing the importance of male reproductive fitness in determining reproductive success and population persistence, the physiological basis of male reproductive maturation remains poorly understood in this mosquito species. Interestingly, our preliminary observation revealed an enriched expression of an LPS-induced TNF-α factor (LITAF) homolog, LL6, in the reproductive tissues of *An. culicifacies* following mating raises the possibility that immune–apoptotic regulators are repurposed to maintain sperm homeostasis. While no direct role of LITAF members in fertility has been reported in vertebrates or invertebrates, its involvement in lysosomal degradation, autophagy, and oxidative stress responses [^26^] allowed us to hypothesize and test whether the identified mosquito LITAF may facilitate testicular homeostasis regulation and phagocytic clearance of defective germ cells, allowing for age-dependent replenishment and preserving male reproductive competence.

To test it, we combine Mix-n-Fix mating assays, microscopic imaging of gonadal development, and functional interrogation of *AcLITAF6* expression through RNAi and cellular assays such as TUNEL staining, sperm viability assessment, and phagocytosis measurement. By linking male age, gonadal maturation, apoptotic germ cells clearance, and the immune–apoptotic role of LITAF/LL6, we propose a dual-layered mechanism involving complement-like clearance coupled with apoptotic signaling that safeguards reproductive fitness in mosquitoes. Our study explores a mechanistic basis for understanding how *An. culicifacies* manages germline turnover, establishing *AcLITAF6* as a potential target for future vector control strategies that exploit the species’ unique reproductive biology.

## Materials and methods

### Mosquito rearing and maintenance

*Anopheles culicifacies* (sibling species A) mosquitoes were reared and maintained in the central insectary of the National Institute of Malaria Research under controlled environmental conditions of 28 ± 2 °C, 60–80% relative humidity, and a 12:12 h light/dark cycle. Eggs were placed on the surface of deionized water and allowed to hatch. Larvae were reared at a density of approximately 1000 individuals per enamel tray (35 cm diameter × 15 cm depth) containing water supplemented with a mixture of fish food (Gold Tokyo, India) and dog food (Pet Lover’s Crunch Milk Biscuit, India) once a day. Pupae were collected in plastic containers using a glass dropper. For emergence, plastic containers containing pupae were transferred to rearing cages (30 × 30 × 30 cm) and maintained on 10% sucrose solution provided via cotton swabs. Female mosquitoes were blood-fed on rabbits to facilitate egg development. Colony maintenance and mosquito rearing were conducted in accordance with the guidelines of the Institutional Animal Ethics Committee of NIMR (Approval No. NIMR/IAEC/2017-1/07) [^22^].

### Morphometric assessment of reproductive organs

For the morphological study, mosquitoes of both sexes of different ages, ranging from just emerged to 13 days old, were anesthetized and placed in a drop of DEPC-treated water on a dissecting slide. Then, the terminal abdominal segment was stretched using a sharp needle to dissect the reproductive tissue and visualized under the camera-fitted microscope (Magnus MLX plus).

The sperm reservoir region within the testes was identified based on a consistent morphological criterion. Specifically, the reservoir was defined as the compact testicular region enriched with mature spermatozoa, characterized by a dense thread-like appearance that was readily distinguishable from the upstream spermatocyst-containing region. The boundary of the sperm reservoir was determined by the clear morphological transition between these two regions. Sperm reservoir size was quantified by measuring the length of the testis occupied by the morphologically distinct sperm-containing region. Measurements were performed on dissected testes using microscopy images and were based on criteria previously described for reproductive morphology analyses in *Anopheles* mosquitoes. The accessory gland-associated clear area was identified as the translucent region surrounding and immediately adjacent to the male accessory glands (MAGs) secretary cells, lacking the dense cellular appearance [^27^].

For comparative evaluations, the captured images of both male reproductive organ (MRO) and female reproductive organ (FRO) at 4X, 10X, and 40X magnifications were imported, and all relevant parameters were calibrated under an isogenic environment of the automated software (Magnus Analytics MagVision, version-x64).

### Mating status and Mix-n-fix experiments

Under a stereomicroscope, mosquito pupae were separated into male and female groups based on terminalia morphology and kept in separate plastic containers partially filled with deionized water to maintain virgin males and females. For adult emergence, these pupae-containing containers were placed in rearing cages (30 × 30 × 30 cm) with cotton pads soaked in 10% sucrose solution. Cages were examined after adult emergence to ensure that all mosquitoes were of the same sex.

For mix-n-fix mating experiment, male mosquitoes of various ages (0-24 hrs to 9-10 days old) were mixed with a female of a fixed age range (3–4 days) in a ratio of 2:1 (100 males & 50 female mosquitoes) for the mating experiment and vice -versa i.e. a fixed age of male (4-5 days old) mixed with various age female.

For mating assays, 3–4-day-old virgin females were introduced overnight into cages containing an equal number of age-matched virgin males (100 males and 100 females per cage). Mating success was verified the following morning by dissecting a random subset of females (n= 4-5) and examining spermathecae for motile sperm under a light microscope. After 48 hours of mixing, female mosquitoes that had been starved for two to six hours were allowed to feed on rabbits for two hours to facilitate blood meal acquisition. To distinguish fully fed mosquitoes from unfed or partially fed mosquitoes, their abdomen was carefully examined.

### In silico analysis: Domain arrangement and structural modelling of Anopheles culicifacies LITAF6 (AcLITAF6)

The putative *AcLITAF6* gene was retrieved from the hemocyte RNA-Seq data of naïve *An. culicifacies* [^24^]. Initial BLASTx analysis against the NCBI NR database yielded significant hits to LITAF-like proteins from multiple mosquito and insect species. To retrieve a full-length transcript, BLASTn analysis was performed against the genome-predicted transcript database of the *An. culicifacies* mosquito, which is available on www.vectorbase.org. Top 10–15 blast hits FASTA sequences were selected from mosquito and non-mosquito species, followed by alignment using ClustalW for multiple sequence alignment analysis [^28^]. The phylogenetic tree was generated from an aligned ClustalW file as input to the maximum-likelihood program of the MEGAX software. The program was run on 1000 bootstraps to confer branching reliability [^29^]. The three-dimensional structure of the LITAF protein was modelled using the I-TASSER server [^30^], and the structure was validated by examining the Ramachandran Plot, which was constructed using RamPlot (Fig. S3 b) [^31^]. For interaction analysis of LITAF with phosphoethanolamine (PE), both the 3D structures of LITAF (with and without zinc) were considered. Docking was performed using Autodock Vina [^32^].

#### Mosquito tissue sample collection

Adult mosquitoes (3-4 days old) were anesthetized by placing them at 4°C for 4-5 minutes before tissue collection. Tissues such as midgut, ovary, spermatheca, testes, male accessory glands, carcass, and the entire male reproductive tract were dissected under a stereomicroscope in DEPC-treated nuclease-free water to reduce RNA degradation on a sterilized slide, following standard procedures. In addition, different developmental stages—eggs, four larval instars, pupae, and adults were collected separately. All tissues were collected in TRIzol reagent for RNA extraction and stored at -80°C till RNA extraction.

### Total RNA isolation and cDNA synthesis

Total RNA was extracted from dissected tissues (midgut, ovary, spermatheca, testes, male accessory glands, carcass, and whole male reproductive tract) and developmental stages (eggs, four larval instars, pupae, adults) of *An. culicifacies* using TRIzol Reagent (RNAiso Plus, Takara Bio, Kusatsu, Japan), following the manufacturer’s instructions [^23^]. RNA quality and concentration were assessed using a NanoDrop 2000c spectrophotometer (Thermo Scientific, USA). Total RNA (∼1 µg) was reverse transcribed into first-strand cDNA using the PrimeScript™ 1st Strand cDNA Synthesis Kit (Takara Bio, Japan). The integrity of the synthesized cDNA was verified by amplification of the actin gene, which served as an internal reference as described earlier [^33^].

### Gene expression analysis

Differential expression of selected transcripts was assessed using quantitative real-time PCR (qPCR) gene expression analysis. Sequenced cDNA or vector-based extracted sequences were used to design primers (https://primer3plus.com/; Supplementary Table 1). Relative expression of chosen target genes in various biological conditions was determined using the SYBR green qPCR master mix and a Bio-Rad real-time PCR machine. All qPCR protocols were identical for each pair of primers with an initial denaturation step at 95°C for 5 min, followed by 40 cycles of 10 sec at 95°C, 15 sec at 52°C, and 22 sec at 72°C. The fluorescence reading was taken at 72°C after each cycle. The latter stages were completing PCR at 95°C for 15 seconds, then switching to 55°C for 15 seconds, and then back to 95°C for 15 seconds, all before creating a melting curve [^25^].

The above protocol was followed with three independent biological replicates for each experiment. Based on expression stability analysis relative to the candidate reference genes, viz. RpS7 and actin; the actin was selected as the internal control for data normalization and analyzed by the 2-ΔΔCt relative quantification method. Finally, GraphPad Prism 11.0.0 was used to generate relative expression graphs. The significance of the treatment was assessed using a one-way ANOVA followed by multiple comparisons.

### dsRNA-mediated gene knockdown

For target gene knockdown (*AcLITAF6*), gene-specific dsRNA primers carrying a T7 promoter overhang were used to amplify complementary DNA (cDNA) by PCR (Supplementary Table S1). Quantification of the purified PCR result was done using a Nanodrop 2000 spectrophotometer (Thermo Scientific). Gene JET PCR purification Kit (Thermo Scientific, Cat #K0701) was used for purification of amplified cDNA, followed by validation (agarose gel electrophoresis). Subsequently, dsRNA was synthesized using Transcript Aid T7 high-yield transcription kit (Cat# K044, Ambion, USA).

Following purification, 69 nl of purified dsRNA (∼2.3 µg/µl) was injected into the thorax of cold-anesthetized, 1–2-day-old adult male mosquitoes using a Nano-injector (Drummond Scientific, CA, USA). For the control group, age-matched mosquitoes were injected with *dsGFP* bacterial-origin dsRNA. Three days after dsRNA injection, target tissues (fat body and male reproductive organs) were dissected from 20–25 mosquitoes, and gene silencing efficiency was assessed using quantitative PCR (qPCR) using both experimental and control groups, as described above.

For video recording, male reproductive tracts were dissected in 1× PBS and mounted under a coverslip. Sperm were released by gently tapping the coverslip with a fine needle and were immediately imaged using a camera-fitted compound microscope. Video acquisition was initiated immediately after sperm release to minimize alterations in sperm viability and motility resulting from extended exposure outside the reproductive tract.

### Assessment of ovary development

For the ovary assessment assay, female mosquitoes from both the control and knockdown groupwere provided a rabbit blood meal. Only fully engorged females were selected for further experiments, while partially fed or unfed mosquitoes were excluded. Approximately 60 hours post-blood meal, mosquitoes were anesthetized, and ovaries were dissected in 1X phosphate-buffered saline (PBS) under a binocular microscope. The number of mature oocytes per ovar**y** was counted manually and compared across experimental groups.

### Terminal deoxynucleotidyl transferase dUTP Nick End Labeling (TUNEL) assay

Apoptotic activity within the male reproductive organ (MRO) was assessed using the Invitrogen™ Click-iT™ Plus TUNEL Assay Kit. Male reproductive organs were dissected in 1× phosphate-buffered saline (PBS) and fixed in 4% paraformaldehyde for 15 min at room temperature. Following fixation, tissues were washed three times with PBS and permeabilized with 0.25% Triton X-100 in PBS for 20 min. Samples were then processed according to the manufacturer’s instructions for the TUNEL staining kit. After labeling, tissues were mounted on glass slides. Fluorescence images were acquired using a confocal microscope (Leica SP8) under identical imaging settings for all samples. TUNEL-positive signals (Alexa Fluor™ 488) were excited with a 488 nm laser and detected between 500–550 nm. TUNEL-positive fluorescence within the reproductive tissues was quantified using ImageJ software and normalized to the tissue area examined [^34^].

### Live/dead sperm viability assay

Viability within the testes was assessed using the LIVE/DEAD™ Sperm Viability Kit (Invitrogen; Thermo Fisher Scientific, #L7011). Testes were dissected from mature virgin control and silenced male mosquitoes in 1× phosphate-buffered saline (PBS) and incubated with SYBR™ 14 working solution (1:50 dilution in DMSO) and Propidium Iodide (PI) according to the manufacturer’s instructions for 2–3 min in the dark at room temperature. Following staining, the testes were mounted on glass slides for imaging [^17^].

Fluorescence imaging was performed using a Leica SP8 confocal microscope under identical acquisition settings for all samples. SYBR™ 14-positive viable sperm were excited with a 488 nm laser and detected between 500–550 nm, whereas PI-positive non-viable sperm were excited with a 561 nm laser and detected between 580–650 nm. Images were processed and analyzed using ImageJ software, and the relative abundance of live and dead sperm within the testes was quantified from multiple biological replicates.

### *In-vivo* phagocytosis assay

For the phagocytosis assay, 48hrs post dsRNA injection into male mosquitoes (1-2 days), as previously mentioned, 69 nl of pHrodoTM Green E. coli BioParticles^TM^ Conjugate (ThermoScientific, Catalogue # P35366) were injected into the hemocoel through the thorax [^35^]. After injection, the mosquitoes were allowed to recuperate and resume physiological functions. To facilitate visualization, sterilized surgical blades were used to remove the wings and legs of mosquitoes. Then, the mosquitoes were gently squished between the coverslip and slide for imaging. A confocal microscope (Olympus FLUOVIEW FV1000) was used to take the pictures. In the control group, 1xPBS-injected mosquitoes received a bioparticle conjugate injection and were visualized as described for silenced mosquitoes.

### Statistical analysis

Statistical analyses were performed using one-way ANOVA for multi-group comparisons, followed by Dunnett’s post-hoc test using the control group as reference. Age-dependent datasets were analyzed by repeated-measures ANOVA with Dunnett’s correction for multiple comparisons. Pairwise comparisons between control and gene-silenced groups were evaluated using an unpaired two-tailed Student’s *t*-test. Statistical significance was set at *p* < 0.05, and exact *p*-values are provided in the figure legends. Additionally, 95% confidence intervals (CI) for mean differences were also calculated.

## Results

### Male fertility follows a non-linear age-dependent pattern

Our preliminary assessment indicated that newly emerged (<24 h old) males of *Anopheles culicifacies* may rapidly attain functional sexual maturity, as observed with proper genitalia rotation, successful mating, sperm transfer to the female spermatheca, and subsequent egg laying by mated gravid females (Supplementary Fig. S1). This observation prompted us to investigate the influence of male age on reproductive output. We therefore performed Mix-n-Fix mating assays in which the age of one sex was held constant while the age of the opposite sex was systematically varied (Fig. 1A). Female egg production following mating was used as an indirect measure of male reproductive performance.

**Fig 1.**
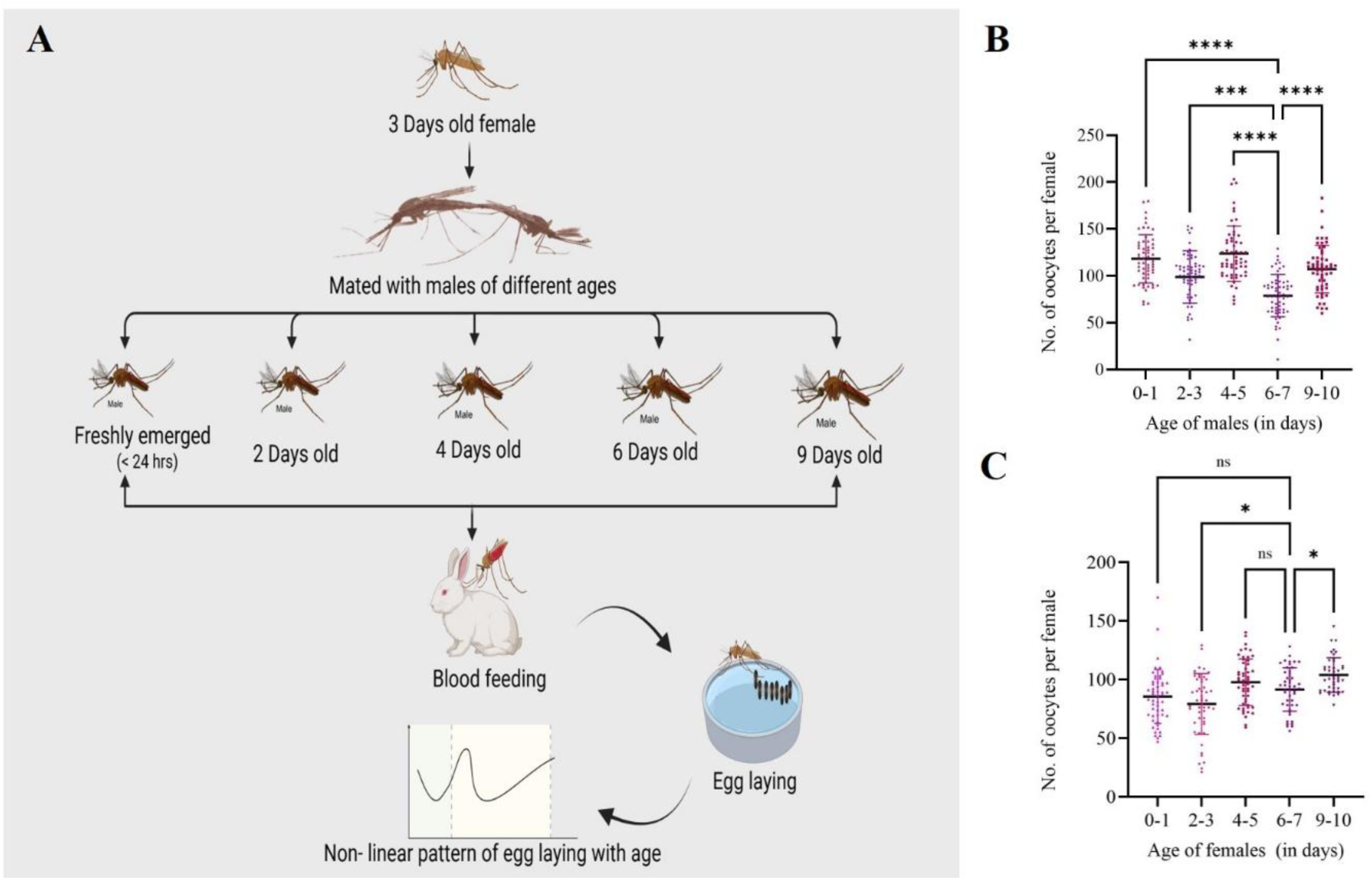
Sex-specific effects of age on reproductive output in *An. culicifacies*. (A) Schematic representation of the Mix-and-Fix mating assay. (B) Quantification of female oocyte numbers after mating with males of different ages (n = 20 females per group, N = 3 independent biological replicates). (C) Quantification of oocyte numbers in females of varying ages after mating with fixed-age males (n = 20 females per group, N = 3 independent biological replicates). Statistical analysis was performed using one-way ANOVA followed by Dunnett’s multiple-comparison test using the 6–7-day-old female group as the reference control. Significance levels are indicated as ns= not significant; P=0.014; P = 0.019; P= 0.0002; ****P < 0.0001.

Interestingly, we observed a non-linear pattern of fertility in both sexes. Male reproductive maturity peaked at 4–5 days of age (Fig. 1B), while female fertility remained relatively stable from fresh emergence to 4 days of age, with no significant age-dependent variation observed during this period (Fig. 1C). These results indicate a bi-phasic regulation of sperm replenishment and fertility, rather than a monotonic decline with age [^11^].

### Testicular development is synchronized with reproductive potential

To explore a possible correlation with a non-linear pattern, we next examined morphological changes with age in the male reproductive organ. Microscopic imaging revealed age-dependent dynamics in testicular architecture, including changes in spermatocyst number, sperm reservoir size, and accessory gland morphology [^36^]. Newly emerged males contained 0–2 spermatocysts, rising to 3–5 by day 1, but declining progressively thereafter, with no spermatocysts detected in 11–13-day-old virgin males (Supplementary Fig. S2). By contrast, the sperm reservoir area relative to total testes length exhibited dynamic age-dependent changes, eventually occupying nearly the entire testis length by day 13 (Fig. 2A & B). These nonlinear patterns in gonadal morphology closely paralleled the fertility outcomes observed in behavioral assays. Accessory gland morphology also showed age-related changes, with a clear area surrounding the glands evident in 0–1-day-old males but lost by day 3 in virgins, though retained longer in some mated males (Fig. 2C& D). In pioneering work in male *An. gambiae* observed that the morphological character of male accessory glands (MAG), i.e., the presence of a clear area around them, decreased with age in virgin males [^27^]. These morphological changes mirrored the non-linear fertility patterns observed in mating assays [^7^].

**Fig 2.**
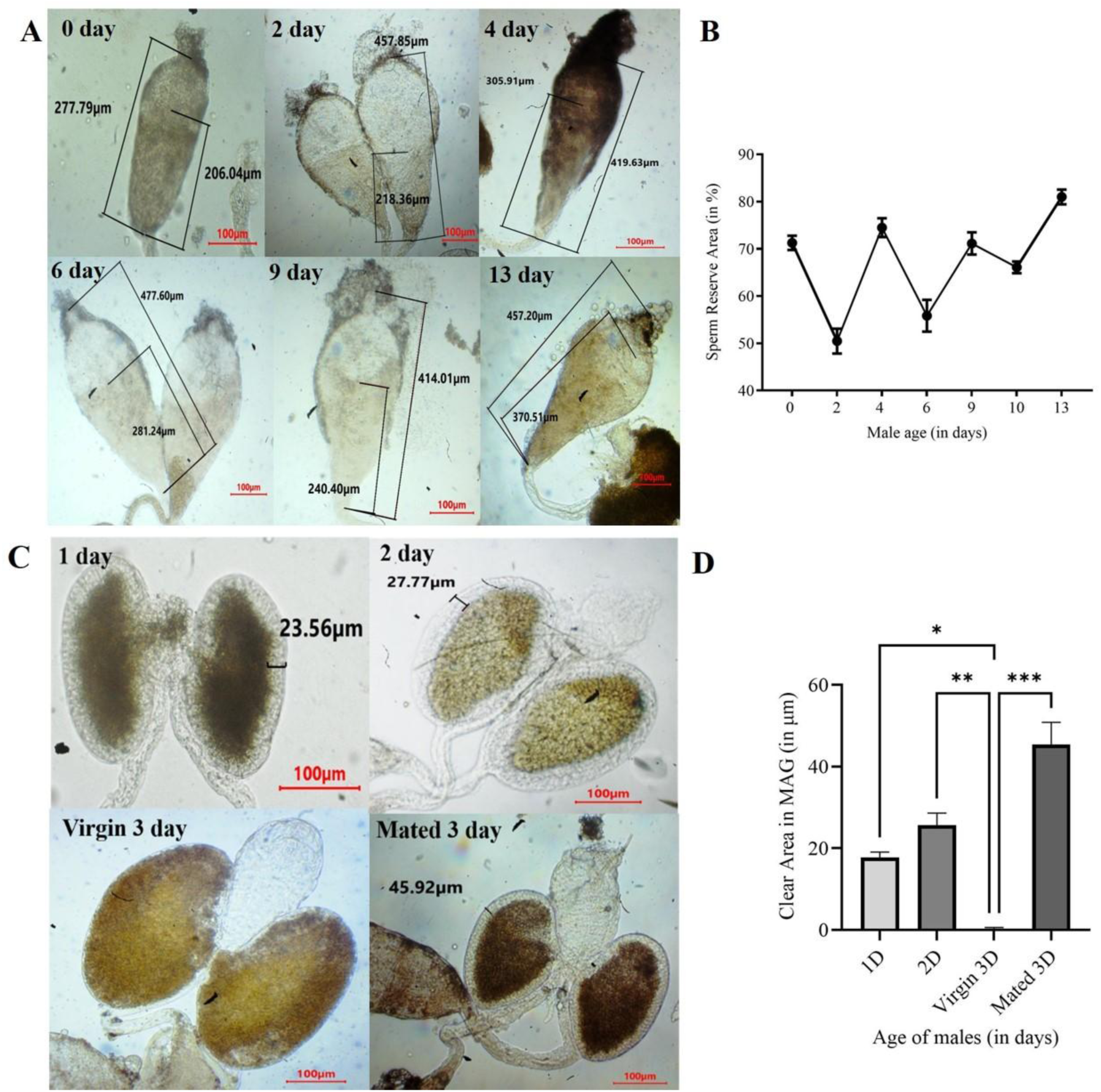
Microscopic imaging and morphometric analysis of *An. culicifacies* testes and male accessory glands (MAGs). (A) Representative microscopic images (10X) of testes from virgin males at sequential ages. (B) Morphometric quantification of sperm reservoir area relative to total testes size across age groups. (C) Representative microscopic images (10X) of male accessory glands (MAGs) from virgin males aged 1–3 days and mated 3-day-old males. (D) Morphometric quantitative analysis of the clear area surrounding the MAGs during age progression and after mating. Data are presented as mean (n = 5 testes per group, N = 3 independent biological replicates). Statistical analysis was performed using one-way ANOVA followed by Dunnett’s multiple-comparison test. For panel B and D, the 2^nd^ day male and 3^rd^ day virgin male are used as references, respectively. Significance levels are denoted as ns=not significant; P =0.011; P = 0.003; P = 0.0003; ****P < 0.0001.

### *In-silico* characterization predicts *AcLITAF6* role in mosquito reproductive physiology

Homology search analysis of an 885 bp long transcript originally identified from hemocyte RNAseq data [^24^], encoding for LITAF-6-like protein, and its enriched expression in reproductive organs, than other organs, viz., head, midgut, and carcass in female mosquitoes (Supplementary Fig. S3), indicated its role in mosquito reproductive physiology. A comprehensive *in silico* analysis against *An. culicifacies* database (www.vectorbase.org) further reveals the identity of the full-length gene (1525 bp) consisting of one intron (1144 bp) spanning between Exon-1 (275 bp) and Exon-2 (106 bp), totaling a 381-bp full-length transcript, here named *AcLITAF6* (ACUA022039). This gene codes for a 126-aa polypeptide (Fig. 3A; Supplementary Fig. S4).

**Fig 3.**
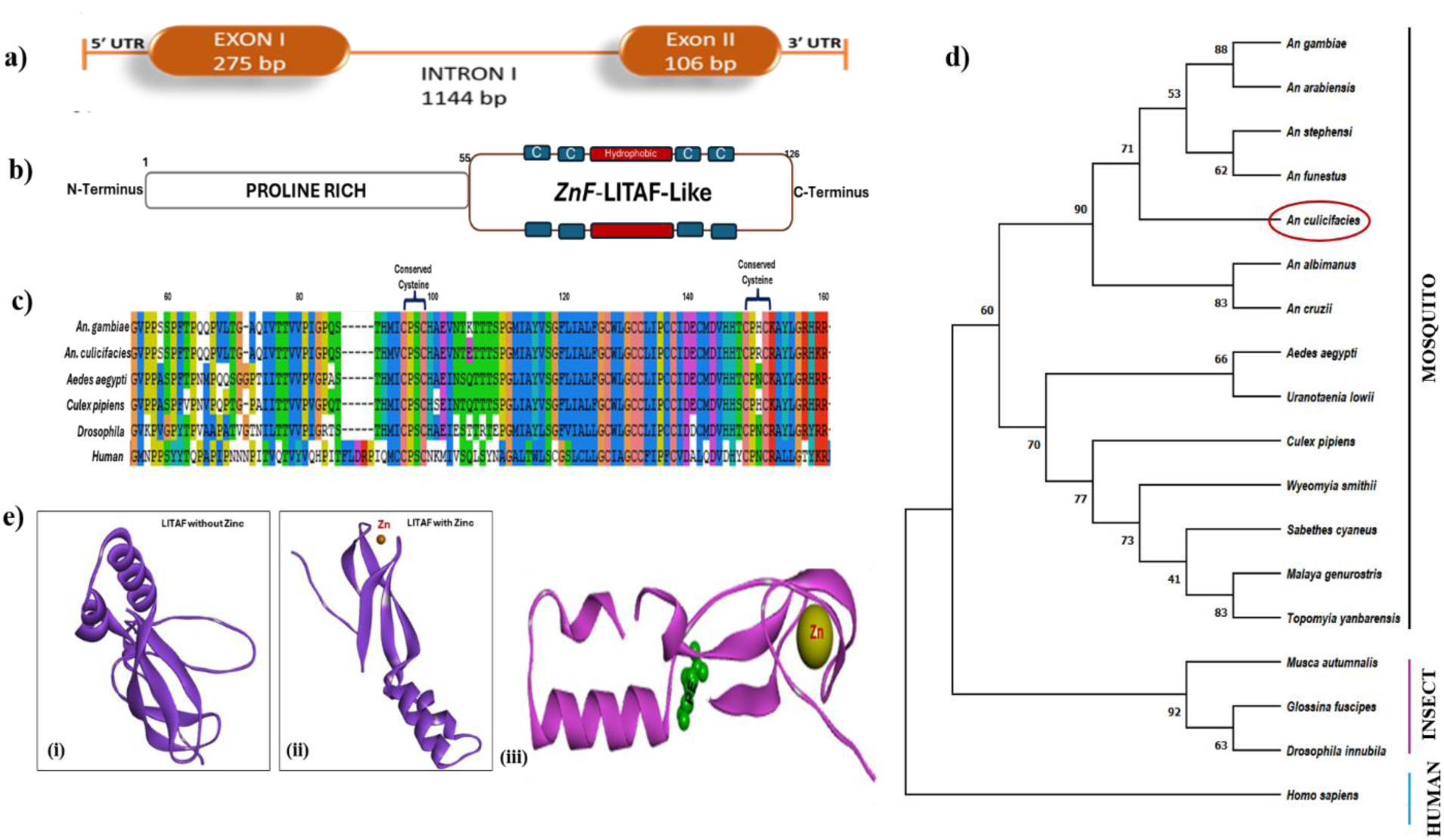
Sequence and structural analysis of *AcLITAF6*. (A) Genomic organization of the *AcLITAF6* transcript showing exon–intron architecture. (B) Domain annotation showing conserved zf-LITAF domain and structural features. (C) Snapshot of multiple sequence alignment of mosquitoes and insects with known human highlighting conserved cysteine residues, a characteristic feature of Zn-LITAF-like proteins encoded by insects. (D) Maximum-likelihood phylogenetic tree generated using MEGA X with 1000 bootstrap replicates. (E) Predicted three-dimensional protein models of *AcLITAF6*, showing strong binding to Zn (i-without; ii-with Zn ion) and PE (iii-Zn and PE/Phosphoethanolamine: PE marked in green).

Pfam-based domain prediction of the encoded ∼14 kDa protein highlighted that it contains a zf-LITAF-like functional domain at 55-126 amino acid residues. The N-terminal contains proline-rich residues likely to support protein-protein interactions, while the C-terminal region, which encodes ‘LITAF domain’, comprises conserved cysteine residues separated by a hydrophobic region (Fig. 3B). Multiple sequence alignment of selected LITAF homologues from diverse orders of insects, *AcLITAF6* showed the presence of sequence conservation at the N-terminal of the protein (Fig. 3C; Supplementary Fig. S5), and a phylogenetic analysis of selected sequenced revealed formation of multiple clades among which *An. culicifacies* found closely related to blood feeder mosquitoes (Fig. 3D). Molecular docking assay revealed conserved structural motifs and predicted binding to Zn²⁺ and phosphatidylethanolamine (PE) (Fig. 3E; Supplementary Fig. S6), comparable to the binding patterns reported in other systems [^37^]. The LITAF protein contains a conserved cysteine-rich domain that coordinates zinc ions (Zn²⁺), which are essential for maintaining its structural stability and proper folding. In addition to zinc binding, the LITAF domain also interacts specifically with phosphatidylethanolamine (PE), a major phospholipid component of cellular membranes. These interactions facilitate membrane localization and functional activity of the protein. Because phospholipid-rich membrane domains are central to apoptotic cell recognition and engulfment processes, the observed PE interaction suggests a potential role for LITAF in membrane-associated cellular clearance pathways such as gamete turnover and debris removal, within the reproductive organs.

### *AcLITAF6* expression and RNAi-mediated silencing reveal its essential role in testicular homeostasis and male fertility

Compared to larvae, the expression profiling of *AcLITAF6* across developmental stages revealed strong upregulation in eggs (Fig. 4A; Supplementary Fig. S7), and male pupae than female (Supplementary Fig. S8). This expression pattern suggests that *AcLITAF6* may contribute to the regulation of apoptotic processes linked to key developmental transitions, particularly during embryogenesis and sex-specific pupal remodelling. Tissue-specific analysis demonstrated that *AcLITAF6* is also highly expressed in the reproductive organs (Fig 4B) of virgin males compared to mated males (Fig 4C), supporting its potential role in regulating male fertility in response to mating. Further observation revealed enriched expression in the testes than the male accessory glands (Fig 4D), pointing to the major function of *AcLITAF6* in sperm production in testes rather than seminal fluid production. These patterns suggest that *AcLITAF*6 may likely contribute to a shared immuno-reproductive function in mosquitoes, linking immune signaling with sperm turnover and reproductive tissue integrity.

**Fig 4.**
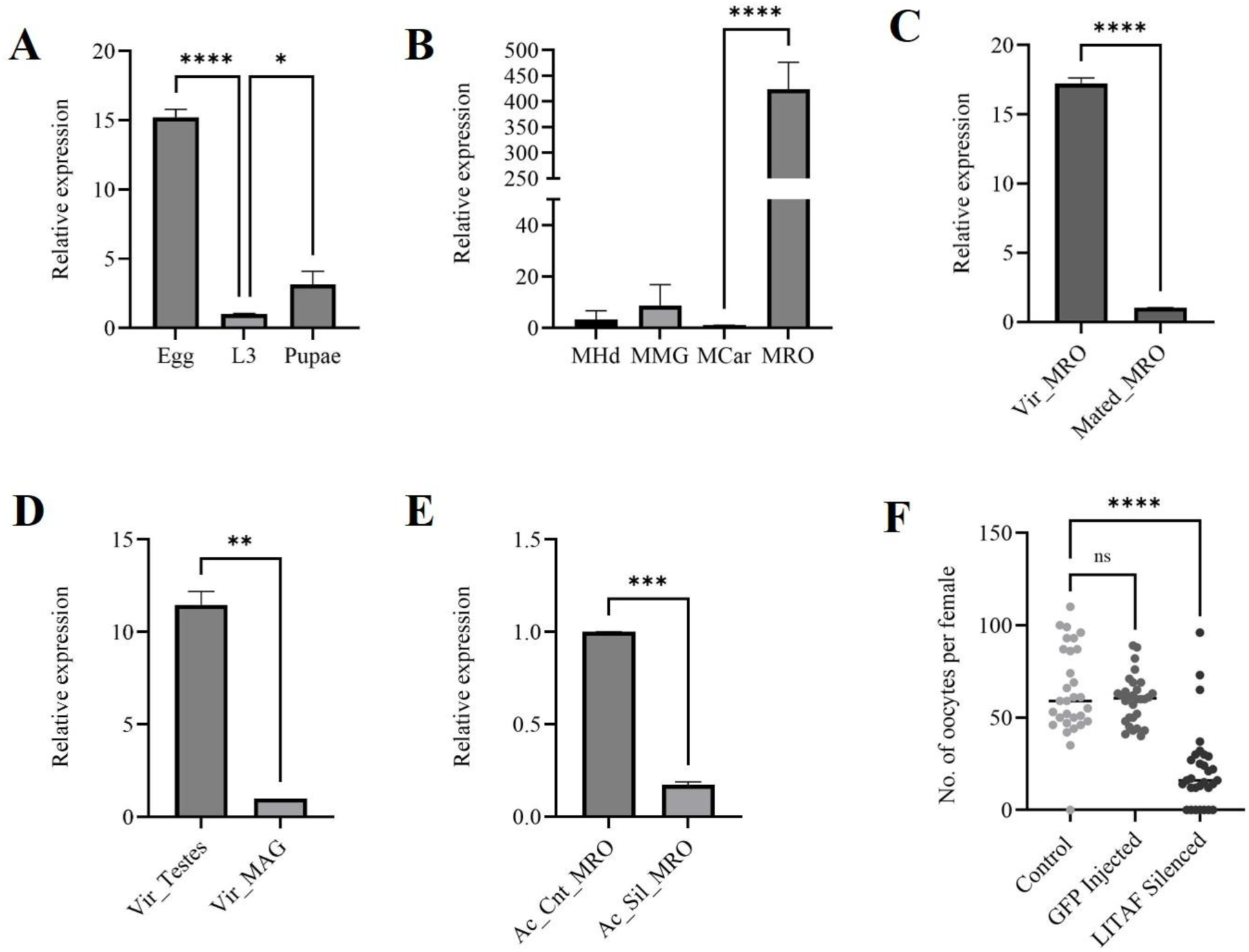
Spatial-temporal expression profiling of *AcLITAF6* in the mosquito *An. culicifacies*. (A) Relative Gene expression across mosquito developmental stages. Eggs (n = 50), larvae (n = 10), pupae (n = 5), and adults (n = 5), N=3. The statistical significance relative to the 3^rd^ instar larva (L3), used as the control, is denoted as P = 0.016, and ****P<0.0001 using one-way ANOVA with Dunnett’s multiple comparisons test. (B) Tissue-specific expression analysis of *AcLITAF6* in 3–4-day-old adult males (n=25, N=3). The statistical significance relative to the male carcass (MCar), used as the control, is denoted as ****P<0.0001 using one-way ANOVA with Dunnett’s multiple comparisons test. (C) Comparative expression of *AcLITAF6* in virgin and mated male reproductive organs (n = 20, N = 3). The statistical significance relative to the mated MRO, used as the control, is denoted as ****P<0.0001 using Unpaired Two-tailed Student’s t-test. (D) Relative expression of virgin testes and male accessory glands (MAG) (n = 25, N = 3). The statistical significance relative to the virgin MAG, used as the control, is denoted as P = 0.0023 using an unpaired two-tailed Student’s t-test. (E) Knockdown validation of *AcLITAF6* in male reproductive organs (MRO) by qPCR (n = 20, N = 3). The statistical significance relative to the virgin male mosquitoes, used as the control, is denoted as P = 0.0002 using the Unpaired Two-tailed Student’s t-test. (F) Quantification of oocyte numbers in females mated with control, dsGFP-injected, and *AcLITAF6*-silenced males. The statistical significance relative to the virgin male mosquitoes, used as the control, is denoted as ****P<0.0001 using the Unpaired Two-tailed Student’s t-test.

To further validate whether *AcLITAF6* is crucial for sperm quality and testicular homeostasis regulation, *AcLITAF6-silenced* male mosquitoes were crossed with healthy females, and a fertility assay was performed. A significant (p<0.0005) reduction in the oocyte counts, but no significant change in hatchability was observed in the gravid females when mated with silenced male mosquitoes (Fig. 5E-F; Supplementary Fig. S9). While we noticed a progressive accumulation of *AcLITAF6* transcripts in the female fat body and ovary at 48 h post-blood meal (Supplementary Fig. S10), mating with *AcLITAF6-silenced* males not only resulted in abnormal ovarian follicle development, and altered morphology of the laid eggs, also affected male testicular and accessory gland architecture (Supplemental Fig 11). Together, these observations pointed out that LITAF may significantly influence overall reproductive outcome, possibly through debris clearance during fertilization and gametogenesis, though the exact mechanism is yet to be unravelled.

**Fig 5.**
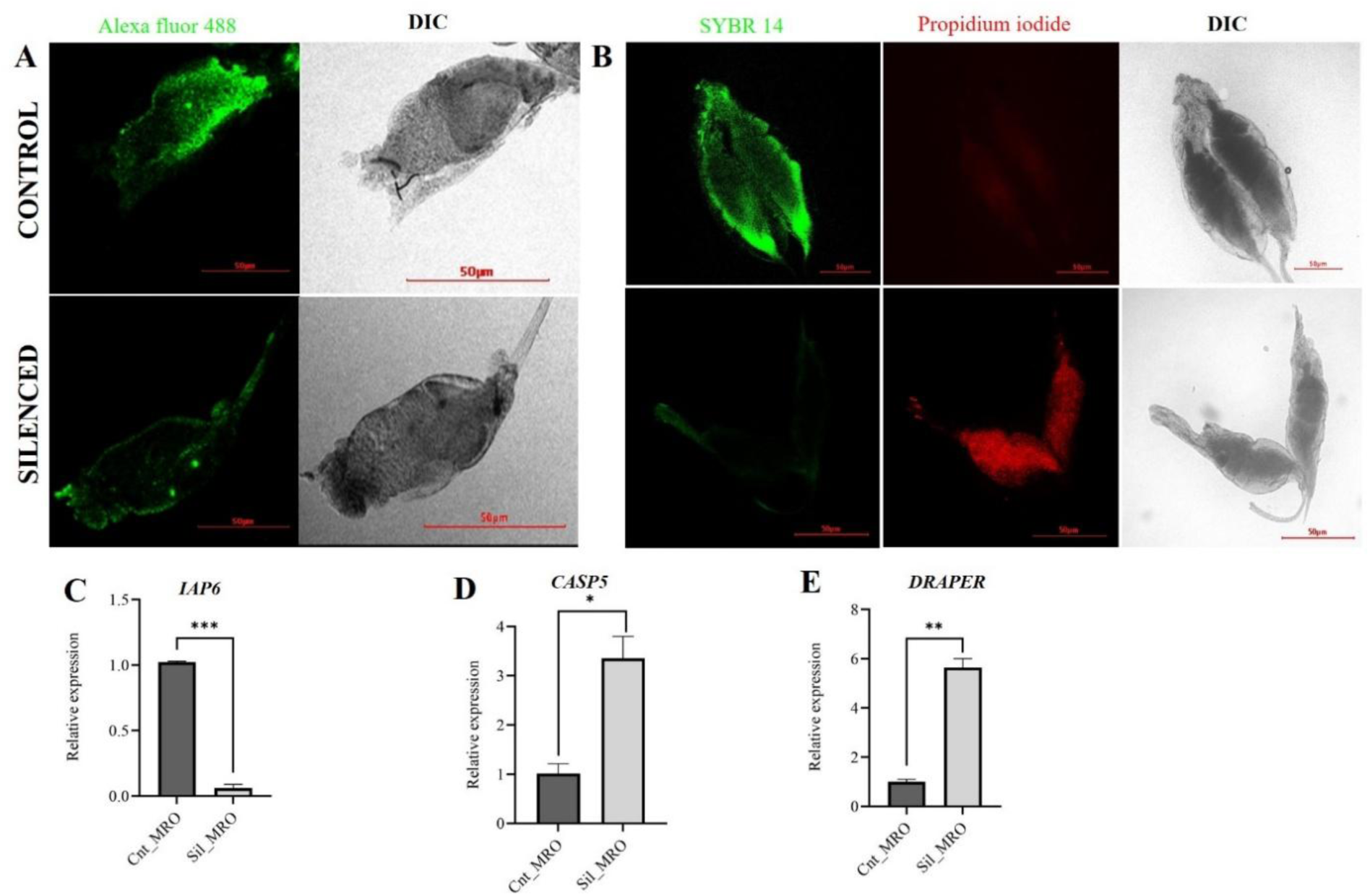
Assessment of *AcLITAF6* silencing on sperm apoptosis and viability. (A) Representative confocal images <u>(10X)</u> of testes from control and *AcLITAF6*-silenced males examined for apoptotic activity via TUNEL assay. Green fluorescence represents TUNEL-positive apoptotic cells. Scale bar = 50 μm. (B) Representative confocal images <u>(10X)</u> of sperm from control and *AcLITAF6*-silenced testes males examined through a live/dead sperm viability assay. Live sperm were stained green with SYBR-14, whereas dead sperm were stained red with propidium iodide (PI). Scale bar = 50 μm. (C-E) Relative expression of genes associated with apoptosis (IAP6 and CASP5) and apoptotic cell clearance/phagocytosis (Draper) was assessed following *AcLITAF6* silencing (n = 20, N = 3). Statistical significance was determined using an unpaired two-tailed Student’s t-test. P = 0.02, P= 0.002, P= 0.0005.

### *AcLITAF6* influences apoptosis-associated testicular homeostasis in the male reproductive organ

To directly assess the role of *AcLITAF6* in maintaining testicular homeostasis, we examined apoptosis-associated cellular activity and sperm viability in the male reproductive organ (MRO). TUNEL staining assay revealed a marked reduction in apoptotic signal in *AcLITAF6* - silenced MROs compared with control males, indicating disruption of normal apoptotic activity within the reproductive tissue (Fig. 5A; Supplemental Fig 12). Consistent with this observation, transcripts associated with apoptotic regulation were quantified. *AcLITAF6* silencing resulted in significant downregulation of inhibitor of apoptosis protein (IAP), accompanied by increased expression of CASPASE5 and Draper, a marker associated with the recognition and clearance of apoptotic cells, which may reflect a compensatory response to impaired cellular homeostasis within the testes. (Fig. 5C-E). Altered expression of these apoptosis-associated markers suggests that depletion of *AcLITAF6* perturbs cellular pathways involved in the regulation and execution of programmed cell death.

Because controlled elimination of defective sperm is essential for maintaining sperm quality, we next examined sperm viability using a live/dead staining assay. *AcLITAF6* -silenced males exhibited a significantly higher proportion of dead spermatozoa than control males (Fig. 5B; Supplemental Fig 13; vedioS1). The accumulation of dead sperm in the reproductive tract indicates impaired sperm quality and suggests that cellular mechanisms responsible for sperm turnover are disrupted in the absence of *AcLITAF6*

### *AcLITAF6* contributes to phagocytosis-associated activity in the male reproductive organ

Because an optimal removal of unused or aged sperms is essential to maintain healthy sperm, next, we investigated cellular clearance activity within the mosquito male reproductive organ (MRO), and therefore, phagocytosis was assessed using pHrodo™ Green E. coli BioParticles™ conjugates. Following particle injection, fluorescence was examined in reproductive tissues of virgin and mated males. Virgin males consistently exhibited stronger fluorescence signals within the MRO than mated males (Fig. 6A-B), indicating higher phagocytosis-associated activity before mating. In contrast, fluorescence intensity was markedly reduced in mated reproductive tissues, suggesting that cellular clearance activity declines following sperm transfer. We hypothesize that the decrease in *AcLITAF6* expression post-mating reflects a physiological transition in the male reproductive system rather than protein transfer. As *AcLITAF6* is integral to the maintenance of germline turnover, it is plausible that its expression is down-regulated once the immediate demand for sperm replenishment is mitigated by the recent ejaculation event.

**Fig 6.**
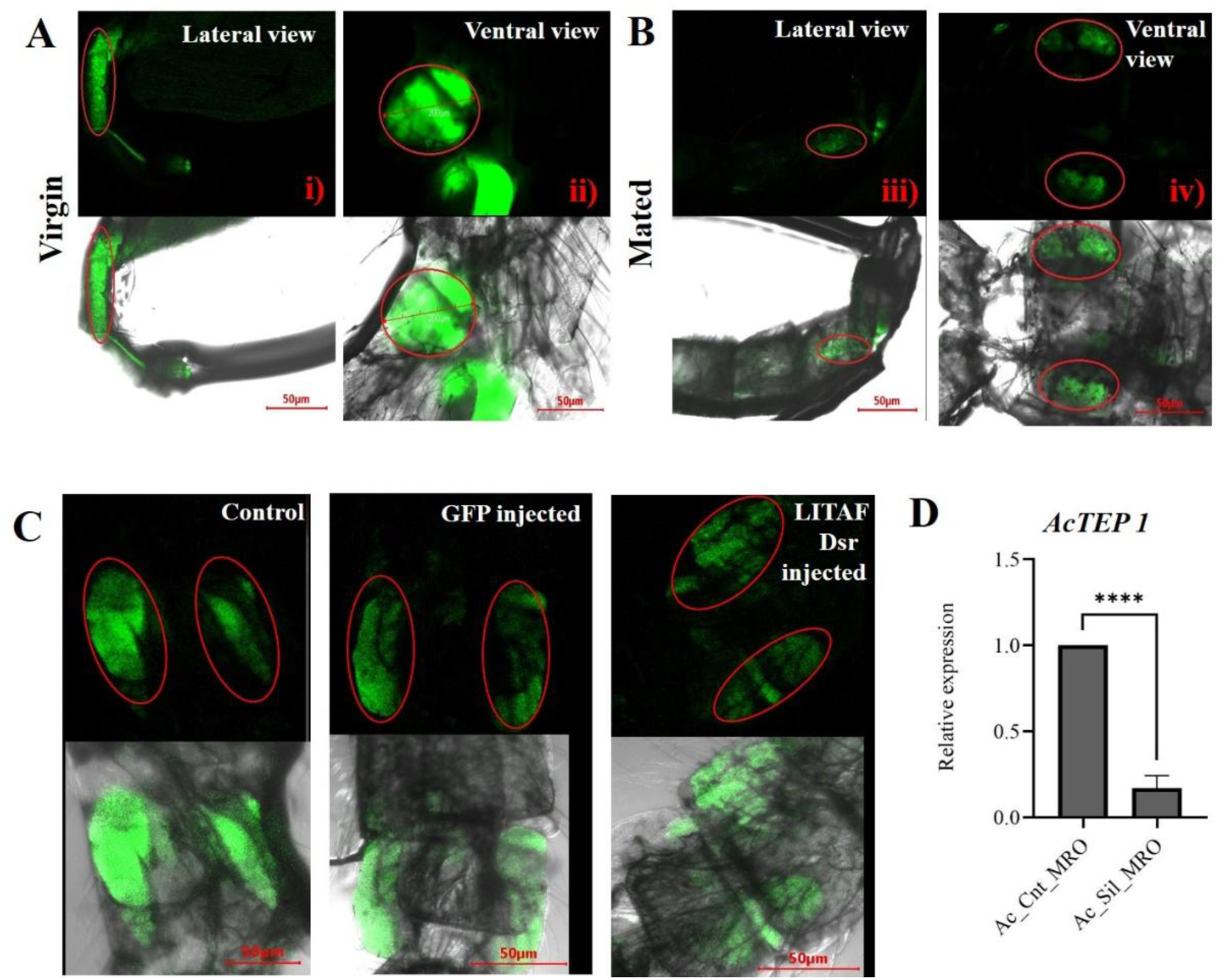
*AcLITAF6* influences phagocytic responses in the male reproductive organ. (A-B) Representative confocal images (10X) of testes in virgin male reproductive organs (MRO) and in mated MRO males injected with pHrodo™ Green bioparticles in adult male *An. culicifacies*. Virgin MRO (Lateral view: i; ventral view: ii) is compared to mated MRO (lateral view: iii; ventral view: iv). Scale bars = 50 μm. (C) Representative confocal images (10X) of testes from control, *dsGFP-*injected, and *AcLITAF6*-silenced males injected with pHrodo™ Green bioparticles to assess phagocytic activity. Scale bars = 50 μm. (D) Relative expression of *AcTEP1* following *AcLITAF6* silencing (n = 20, N = 3). The statistical significance was determined using an unpaired two-tailed Student’s t-test; P < 0.0001.

To determine whether *AcLITAF6* contributes to this process, phagocytic activity was further examined in *AcLITAF6* -silenced virgin males. Silencing of *AcLITAF6* resulted in a pronounced reduction in fluorescence signal compared with control virgin males (Fig. 6C;Supplemental Fig 14), demonstrating impaired uptake or processing of pHrodo-labelled BioParticles within the reproductive organ. Because thioester-containing protein 1 (TEP1) has been implicated in recognition and clearance-associated pathways, its expression was quantified following *AcLITAF6* depletion. TEP1 transcript abundance was significantly reduced in *AcLITAF6*-silenced MROs relative to control tissues (Fig. 6D). The concomitant reduction in pHrodo fluorescence and TEP1 expression suggests that *AcLITAF6* positively regulates cellular clearance mechanisms within the mosquito reproductive tract. Collectively, these findings indicate that the male reproductive organ exhibits active phagocytosis-associated processes that are more pronounced in virgin males and *AcLITAF6* is required for maintaining normal clearance activity within reproductive tissues.

## Discussion

Disclosing the secrets of age-dependent male gonadal changes and assessing their impact on mosquito reproduction could be crucial for designing effective vector suppression strategies [^1^]. It is widely presumed that a gradual decline of fertility may occur with age in male mosquitoes, but conceptually, no evidence exists on this correlation. Surprisingly, our findings reveal that male mosquito fertility is not a simple function of chronological age but follows a non-linear pattern driven by sperm replenishment and gonadal remodelling. Fertility peaks at intermediate ages (4-5 days) correspond to periods of maximal *AcLITAF6* expression and effective germ cells clearance. This challenges the conventional view of a linear decline in fertility and instead suggests a dynamic balance between sperm production, turnover, and the removal of defective germ cells [^9,11^].

In a pioneering work in male *An. gambiae*, it has been observed that the morphological character of male accessory glands (MAG), i.e., the presence of a clear area around them, decreased with age in virgin males [^27^]. Our morphological analyses support this interpretation, showing cyclic remodelling of sperm reservoirs. As spermatocysts decline with age, the reservoir expands, suggesting ongoing sperm release and replenishment. Accessory gland changes further highlight how reproductive tissues adapt to mating status and age, influencing sperm function and fertility outcomes [^6,7^].

While the dynamics of key molecular factors that ensure quality sperm maintenance are still unclear [^17^], suppression of testes expressing the LPS-induced TNF-α factor, i.e., *AcLITAF6*, through RNAi, and subsequent TUNEL and phagocytosis assays demonstrate that *AcLITAF6* is required for the efficient removal of apoptotic germ cells, linking apoptotic signaling with phagocytic clearance. This function parallels mammalian *LITAF6* roles in apoptosis and lysosomal trafficking [^38,39^] and resembles *Drosophila* phagoptosis by cyst cells [^15^]. Together, these results highlight the evolutionary conservation of LITAF-mediated apoptotic regulation and its novel recruitment in mosquito reproductive physiology.

Current studies highlight a complex interplay of male seminal fluid proteins and female physiological responses, including the upregulation of antioxidant genes and active ion regulation, as essential for maintaining sperm viability and optimal female fertility throughout her reproductive lifespan [^40^]. Surprisingly, an observation that mating and blood-feeding boost *AcLITAF6* expression in female reproductive tissues, endorsed its additional role in ovarian remodeling, follicular apoptosis, or immune modulation following insemination and blood-meal digestion [^41,42^]. A reduced reproductive outcome in gravid females that mated with *AcLITAF6*-silenced males supports the idea that LITAF-mediated pathways may serve as a shared immuno-reproductive mechanism in both sexes, orchestrating germline quality regulation and tissue adaptation to reproductive demands.

Modulation of apoptotic marker genes in the LITAF-silenced male mosquito reproductive organ further ensures the integration of complement-like factors (e.g., TEP1) with LITAF-mediated pathways to support a dual-layered system for maintaining sperm quality: immune surveillance to tag defective germ cells and apoptotic-phagocytic clearance to remove them [^43,44^]. This may be crucial for reproductive tissue integrity and fertility across the male lifespan, while in females, *AcLITAF6* may secure optimal ovarian performance after mating and blood-feeding. This sexually dimorphic reproductive behavior in mosquitoes may be intricately tied to molecular regulators like *LITAF6* that integrate immune surveillance with fertility maintenance.

Our findings establish a ’Phagoptotic Surveillance’ model, demonstrating that the *An. culicifacies* testis utilizes evolutionarily conserved immune mechanisms to preserve reproductive fitness through the active clearance of senescent germ cells. Conceptually, the integration of *IAP*, *CASPASE*, and *TEP1* with our structural and functional data supports a framework in which *AcLITAF6* participates in a specialized phagoptotic regulatory mechanism within the mosquito’s male immune-reproductive axis. As this process involves the coordinated recognition and engulfment of damaged or apoptotic cells through membrane-associated signaling, the conserved Phosphatidylethanolamine (PE)-binding property of *AcLITAF6* likely facilitates its recruitment to phospholipid-rich membrane domains involved in apoptotic recognition. This recruitment is functionally equivalent to systemic immune pathways, where *AcLITAF6* acts as a scaffold to integrate apoptotic signaling with phagocytic clearance. Consistent with this, *AcLITAF6* silencing simultaneously altered apoptosis-associated markers, reduced phagocytic activity, suppressed *TEP1* expression, and promoted the accumulation of dead sperm. We propose that *AcLITAF6* functions as a membrane-associated regulator of this immuno-reproductive axis, ensuring testicular homeostasis by preventing the cytotoxic effects of secondary necrosis—a critical quality control checkpoint for maintaining male fertility as the vector ages (Fig. 7).

**Fig. 7.**
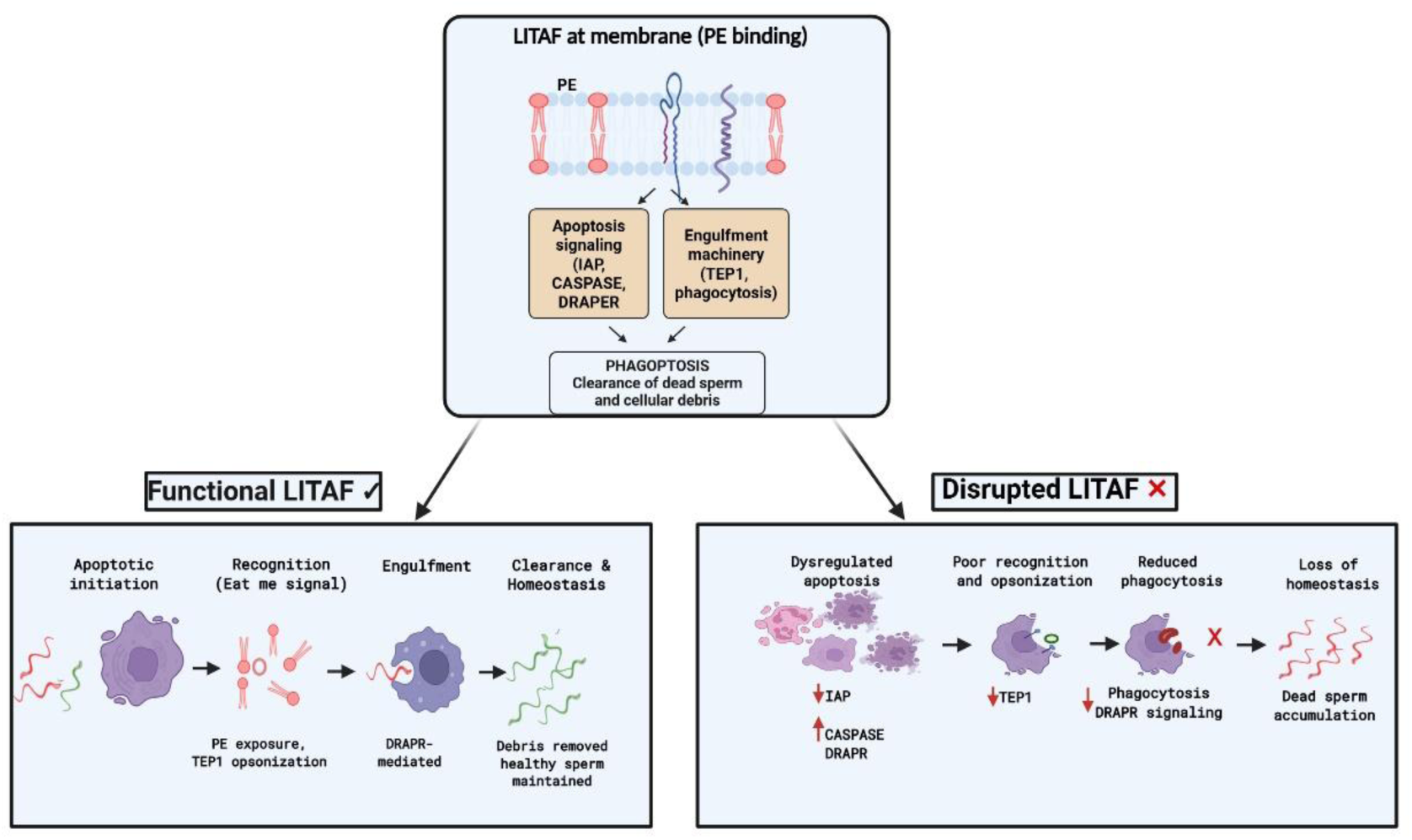
Proposed working model for LITAF function in mosquito immune-reproductive axis: LITAF is predicted to associate with PE-containing membrane domains and coordinate immune-active apoptosis-associated signalling with phagocytic clearance (phagoptosis) in the male reproductive organ. This process promotes the removal of defective sperm and the maintenance of testicular homeostasis. LITAF silencing impairs sperm clearance, increases dead sperm accumulation, and reduces reproductive fitness, resulting in fewer oocytes and abnormal eggs in mated females.

In vector control contexts, targeting such sperm-quality pathways may enhance the effectiveness of genetic techniques by reducing the reproductive competitiveness of aging males. These findings have practical implications for vector control. Understanding the observed non-linear fertility patterns and molecular regulators, such as *AcLITAF6*, provides insight into the optimal timing of male releases and potential molecular targets for fertility manipulation.

## Conclusion

We identify *AcLITAF6* as a novel immuno-reproductive regulator in *Anopheles culicifacies*, linking age, sperm clearance, fertility, and female reproductive adaptation. This discovery advances our understanding of mosquito reproductive biology and highlights new opportunities for integrating molecular and ecological approaches in vector control.

### Limitations and future perspectives

While this study identifies *AcLITAF6* as a pivotal regulator of male fitness in *Anopheles culicifacies*, several limitations remain. Our assessment of male fertility relied on female reproductive outcomes—fecundity and egg morphology—as indirect proxies for ejaculate quality. Future research should incorporate direct measurements of sperm motility and DNA integrity to distinguish between quantitative and functional defects. Furthermore, while *E. coli* bioparticles confirmed a general phagocytic role for *AcLITAF6*, providing definitive proof that testicular homeostasis requires direct visualization of apoptotic germ cells clearance via high-resolution electron microscopy.

Additionally, the observed non-linear fertility patterns would require further validation in wild populations across the species complex. Finally, exploring the independent role of female-derived LITAF in defective eggs will provide a holistic view of this gene’s impact on mosquito population dynamics.

## Supporting information

Supplemental data sheet

## Acknowledgment

We extend our gratitude to the insectary staff for their rigorous efforts in mosquito rearing. We also appreciate Kunwarjeet Singh, Sattey Singh, Lipun, Benudhar Mukhi, and Nishant for their technical support in the laboratory. We are grateful to Dr. Neetu Singh, Technician B, AIRF, JNU, New Delhi, India, for confocal microscopy. Finally, we are highly thankful to ICMR-National Institute of Malaria Research, Delhi, for extending the infrastructural and other logistic support for the Investigation. Work in the laboratory is supported by the Indian Council of Medical Research (ICMR), Government of India. Pooja Rohilla is the recipient of the CSIR Research Fellowship (SRF/09/905(0024)/2020-EMR-I). We thank ICMR (No.6/9-7(287)/2022-ECD-II) for sanctioning a research grant to Rajnikant Dixit. The funders played no role in the design of the study, data collection and analysis, decision to publish, or manuscript preparation.

## Author’s contribution statement

Conceptualization: P.R., J.R., R.D.; Data curation & Formal analysis: P.R., V.S.; Investigation: P.R., V.S., V.Sr., P.Y., N.S., T.S., G.S., J.R.; Methodology: P.R., J.R., G.T., R.D.; Project administration: R.D.; Supervision: R.D., J.R., S.T.; Visualization &Validation: P.R., V.S., J.R.G.T.; Writing – original draft: P.R., V.S.; Writing – review & editing: N.S., T.S., J.R., S.T., R.D. All authors have read and agreed to the published version of the manuscript.

## Data Availability and Declaration

All data generated or analysed during this study are included in this published article [and its supplementary information files].

## Ethical Clearance

Necessary ethical clearance for mosquito colonization was taken from the Institutional Animal Ethics Committee of NIMR (Approval No. NIMR/IAEC/2017-1/07).

## Statement on the Use of Artificial Intelligence (AI)

Artificial intelligence tools were used solely to assist in language refinement and improvement of clarity and coherence of the manuscript. All scientific concepts, interpretations, experimental design, data analysis, and conclusions are entirely the responsibility of the authors. AI was not used to generate, manipulate, or analyse any original data, figures, or results, and all content was critically reviewed and validated by the authors before submission.

## References

1. Knipling EF. Possibilities of insect control or eradication through the use of sexually sterile males. J Econ Entomol. 1955;48(4):459–62.

2. Alphey L, Benedict M, Bellini R, Clark GG, Dame DA, Service MW, et al. Sterile-Insect Methods for Control of Mosquito-Borne Diseases: An Analysis. Vector Borne Zoonotic Dis. 2010;10(3):295–311.

3. Burt A. Heritable strategies for controlling insect vectors of disease. Philos Trans R Soc Lond B Biol Sci. 2014;369(1645):20130432.

4. Helinski MEH, Harrington LC. Male mating biology and mating strategies of mosquitoes. Acta Trop. 2011;121(3):171–8.

5. Hancock PA, Sinkins SP, Godfray HCJ. Strategies for introducing Wolbachia to reduce transmission of mosquito-borne diseases. PLoS Negl Trop Dis. 2016;10(7):e0004874.

6. Clements AN. The Biology of Mosquitoes, Vol. 2. Wallingford: CABI Publishing; 1999.

7. Oliva CF, Damiens D, Benedict MQ. Male reproductive biology of *Aedes* mosquitoes. Parasites Vectors.2014;7:493.

8. Huq F, Farnesi LC, Martins AJ, Valle D. Environmental determinants of mosquito reproductive development. J Insect Physiol. 2010;56(11):1550–7.

9. Charlwood JD, Jones MDR, Hilburn RH, Kihonda J, Billingsley PF. Swarming and mating in *Anopheles gambiae* s.l. Acta Trop. 2002;81(2):101–11.

10. Reinhardt K. Evolutionary consequences of sperm aging. Q Rev Biol. 2007;82(4):375–93.

11. Johnson SL, Gemmell NJ. Are old males still good males? Sperm senescence and fertilization. Biol Lett. 2012;8(4):739–42.

12. Das De, T., Sharma, P., Rawal, C., Kumari, S., Tavetiya, S., Yadav, J., Hasija, Y., & Dixit, R. Sex specific molecular responses of quick-to-court protein in Indian malarial vector Anopheles culicifacies: conflict of mating versus blood feeding behaviour. Heliyon. 2017;3, 361. 10.1016/j.heliyon.2017

13. Radwan J. Male age, sperm competition, and reproductive success. Trends Ecol Evol. 2003;18(10):579–85.

14. Pizzari T, Dean R, Pacey A, Moore H, Bonsall MB. The evolutionary ecology of sperm function: sperm design, quality, and strategic allocation. Philos Trans R Soc Lond B Biol Sci. 2008;363(1505):3169–85.

15. Yacobi-Sharon K, Namdar Y, Arama E. Alternative germ cell death pathway in *Drosophila* involves HtrA2/Omi, lysosomes, and a caspase-independent mechanism. Cell Death Differ. 2013;20(1):130–9.

16. Blandin S, Shiao S-H, Moita LF, Janse CJ, Waters AP, Kafatos FC, Levashina EA. Complement-like protein TEP1 is a determinant of vectorial capacity in the malaria vector *Anopheles gambiae*. Cell. 2004;116(5):661–70.

17. Kumari, S., Tevatiya, S., Rani, J., Das De, T., Chauhan, C., Sharma, P., Sah, R., Singh, S., Pandey, K. C., Pande, V., & Dixit, R. A testis-expressing heme peroxidase HPX12 regulates male fertility in the mosquito Anopheles stephensi. Scientific Reports. 2022; 12(1). 10.1038/s41598-022-06531-x

18. Thailayil J, Magnusson K, Godfray HC, Crisanti A, Catteruccia F. Spermless males elicit large-scale female responses to mating in the malaria mosquito Anopheles gambiae. Proc Natl Acad Sci U S A. 2011 Aug 16;108(33):13677–81. doi: 10.1073/pnas.1104738108. Epub 2011 Aug 8.

19. Yamamoto DS, Sumitani M, Kasashima K, Sezutsu H, Matsuoka H, Kato H. A synthetic male-specific sterilization system using the mammalian pro-apoptotic factor in a malaria vector mosquito. Sci Rep. 2019 Jun 3;9(1):8160. doi: 10.1038/s41598-019-44480-0.

20. Sharma VP, Dev V. Biology & control of Anopheles culicifacies Giles 1901. Indian J Med Res. 2015 May;141(5):525–36. doi: 10.4103/0971-5916.159509.

21. Sharma P, Das De T, Sharma S, Kumar Mishra A, Thomas T, Verma S, Kumari V, Lata S, Singh N, Valecha N, Chand Pandey K, Dixit R. Deep sequencing revealed molecular signature of horizontal gene transfer of plant-like transcripts in the mosquito Anopheles culicifacies: an evolutionary puzzle. F1000Res. 2015 Dec 30;4:1523. doi: 10.12688/f1000research.7534.1.

22. De, T. Das, Thomas, T., Verma, S., Singla, D., Rawal, C., & Srivastava, V. A synergistic transcriptional regulation of olfactory genes derives complex behavioral responses in the mosquito Anopheles culicifacies. Biorivix.2017.

23. Das De T, Sharma P, Tevatiya S, Chauhan C, Kumari S, Yadav P, Singla D, Srivastava V, Rani J, Hasija Y, Pandey KC, Kajla M, Dixit R. Bidirectional Microbiome-Gut-Brain-Axis Communication Influences Metabolic Switch-Associated Responses in the Mosquito *Anopheles culicifacies*. Cells. 2022 May 31;11(11):1798. doi: 10.3390/cells11111798.

24. Rani, J., Chauhan, C., De, T. Das, Kumari, S., Sharma, P., Tevatiya, S., Patel, K., Mishra, A. K., Pandey, K. C., Singh, N., & Dixit, R. Hemocyte RNA-Seq analysis of Indian malarial vectors *Anopheles stephensi* and *Anopheles culicifacies* : From similarities to differencesCAT.Gene.2021;798(June),145810.10.1016/j.gene.2021.14581 <u>0</u>

25. Rani, J., De, T. Das, Chauhan, C., Kumari, S., Sharma, P., Tevatiya, S., Chakraborti, S., Pandey, K. C., Singh, N., & Dixit, R. Functional disruption of transferrin expression alters reproductive physiology in *Anopheles culicifacies*. PLoS ONE.2022; 17(3 March), 1–17. 10.1371/journal.pone.0264523

26. Tang, X., Marciano, D. L., Leeman, S. E., & Amar, S. LPS induces the interaction of a transcription factor, LPS-induced TNF-α factor, and STAT6 (B), with effects on multiple cytokines. Pnas. 2005; 102(14), 2–7. 10.1073/pnas.0501159102

27. Huho, B. J., Ng’habi, K. R., Killeen, G. F., Nkwengulila, G., Knols, B. G. J., & Ferguson, H. M. A reliable morphological method to assess the age of male Anopheles gambiae. Malaria Journal. 2006;5, 1–11. 10.1186/1475-2875-5-62

28. Thompson, J. D., Higgins, D. G., & Gibson, T. J. CLUSTAL W: improving the sensitivity of progressive multiple sequence alignment through sequence weighting, position-specific gap penalties and weight matrix choice.1994; 22(22), 4673–4680.

29. Kumar, S., Stecher, G., Li, M., Knyaz, C., & Tamura, K. MEGA X : Molecular Evolutionary Genetics Analysis across Computing Platforms. 2018; 35(6), 1547–1549. 10.1093/molbev/msy096

30. Lin, Y.F., Cheng, C.W., Shih, C.S., Hwang, J.K., Yu, C.S., and Lu, C.H. MIB: metal ion-binding site prediction and docking server. Journal of chemical information and modelling. 2016;56(12), pp.2287–2291.

31. Kumar, M. and Rathore, R.S. RamPlot: a webserver to draw 2D, 3D and assorted Ramachandran (φ, ψ) maps. Applied Crystallography. 2025; 58(2).

32. Trott, O. and Olson, A.J. AutoDock Vina: improving the speed and accuracy of docking with a new scoring function, efficient optimization, and multithreading. Journal of computational chemistry.2010;31(2), pp.455–461.

33. Saini, V., Rohilla, P., Srivastava, V., Tandon, G., & Yadav, P. Sensory appendage protein triggers alarm to pyrethroid in Indian malarial vector *Anopheles culicifacies*. Plos one. 2025; 1–14. 10.1371/journal.pone.0333483

34. Sharma, P., Sharma, S., Mishra, A. K., Thomas, T., De, T. Das, Rohilla, S. L., Singh, N., Pandey, K. C., Valecha, N., & Dixit, R. Unraveling dual feeding-associated molecular complexity of salivary glands in the mosquito Anopheles culicifacies. Biology Open. 2015; 4(8), 1002–1015. 10.1242/bio.012294

35. Lombardo, F., Ghani, Y., Kafatos, F. C., & Christophides, G. K. Comprehensive Genetic Dissection of the Hemocyte Immune Response in the Malaria Mosquito Anopheles gambiae. 2013;9(1). 10.1371/journal.ppat.1003145

36. Mahmood, F., & Reisen, W. K. *Anopheles culicifacies*: effects of age on the male reproductive system and mating ability of virgin adult mosquitoes. Medical and Veterinary Entomology.1994;8(1), 31–37. 10.1111/j.1365-2915.1994.tb00380.x

37. Ho, A. K., Wagstaff, J. L., Manna, P. T., Wartosch, L., Qamar, S., Garman, E. F., Freund, S. M. V, & Roberts, R. C. The topology, structure and PE interaction of LITAF underpin a Charcot-Marie-Tooth disease type 1C. BMC Biology. 2016; 1–21. 10.1186/s12915-016-0332-8

38. Polyak K, Xia Y, Zweier JL, Kinzler KW, Vogelstein B. A model for p53-induced apoptosis. Nature. 1997;389(6648):300–5.

39. Shin MS, Moghrabi N, Pan J, Lee YH, Pledger WJ. LITAF regulates endosomal/lysosomal trafficking through interaction with the E3 ligase Itch. J Biol Chem. 2010;285(2):1170–9.

40. Baldini, F., Gabrieli, P., South, A., Valim, C., Mancini, F., & Catteruccia, F. The Interaction between a Sexually Transferred Steroid Hormone and a Female Protein Regulates Oogenesis in the Malaria Mosquito Anopheles gambiae.2013; 11(10). 10.1371/journal.pbio.1001695

41. Degner EC, Harrington LC. Mating activity and blood feeding drive reproductive maturation in female *Aedes aegypti* mosquitoes. J Insect Physiol. 2016;93–94:16–23.

42. Camargo, C., Braimah, Y. H. A., Amaro, I. A., Harrington, L. C., Wolfner, M. F., & Avila, F. W. Mating and blood – feeding induce transcriptome changes in the spermathecae of the yellow fever mosquito *Aedes aegypti*. Scientific Reports. 2020;1–13. 10.1038/s41598-020-71904-z

43. Shaw WR, Teodori E, Mitchell SN, Baldini F, Gabrieli P, Rogers DW, Catteruccia F. Mating activates the heme peroxidase HPX15 in the sperm storage organ to ensure fertility in *Anopheles gambiae*. Proc Natl Acad Sci U S A. 2014;111(16):5854–9.

44. Pompon, J., & Levashina, E. A. A New Role of the Mosquito Complement-like Cascade in Male Fertility in Anopheles gambiae. 2015; 1–17. 10.1371/journal.pbio.1002255

