## Supplemental data sheet for "Male Age and Sexual Maturity: Lipopolysaccharide-induced tumor necrosis factor influences sperm quality and reproduction in *Anopheles culicifacies*"

**Table 1. List of primers and their sequences.**

| S. No. | Name of the gene | Left primer | Right primer |
|---|---|---|---|
| 1 | Ac_LPS-induced TNF-α factor 6 *(AcLITAF6)* | GGGCACAGATTGTTACCAC | GTGGACAGGTATGATGGATA |
| 2 | LITAF6_DSR | TAATACGACTCACTATAGGGATGTCGAAAGATGGACCTC | TAATACGACTCACTATAGGGCTAACCAGCATCCAAACAGT |
| 3 | GFP_DSR | TAATACGACTCACTATAGACGACGGCAACTACAAGACC | TAATACGACTCACTATAGGAACTCCAGCAGGACCATGT |
| 4 | Ac_Actin | GCGGTATACTGACACTCAA | CAAACATGATCTGTGTCATC |
| 5 | Ac_RSP7 | ATCGCTATGGTGTTCGGTTC | TTGTTGAACTCGACCTCACG |
| 6 | Ac_Caspase 5 | TCACAGGGCAGATCATCGTA | AATGTGGCGTTCGAAGAAGT |
| 7 | Ac_IAP6 | AAACCTGCCCCTTTTCATCT | CATCGGGCATGAAATCTTTT |
| 8 | Ac_Draper | TCTTTACGATGCGTTGCTTG | AGTGGATA CCATCCCCATCA |
| 9 | Ac_TEP 1 | CCAAGTTGTTGGAGATCAAT | ATTCTGATCGACAACGTACC |

**Fig. S1.**

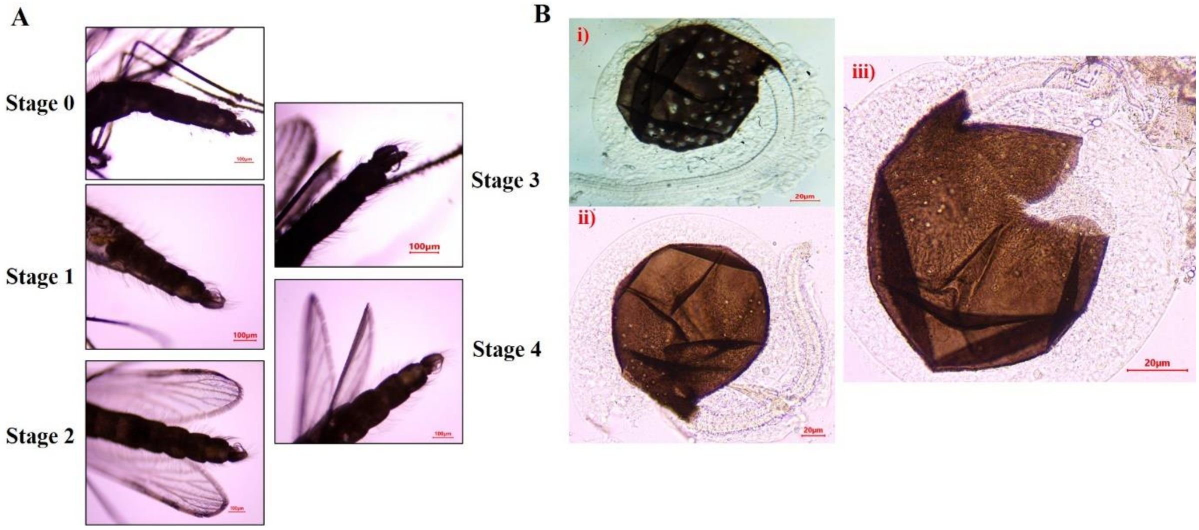


**Fig. S1. Genitalia rotation in male and spermathecal morphology in female mosquitoes.** (A) Representative images showing progressive genitalia rotation during adult male maturation. Stage 0 represents a newly emerged male with unrotated terminalia, while Stages 1–4 depict sequential rotation of the genitalia until complete rotation is achieved. Scale bars, 100 µm.

(B) Representative images of spermathecae from virgin and mated females. (i) Spermatheca of a virgin female showing the absence of stored sperm. Spermathecae of mated females intact (ii) and (iii) ruptured containing sperm bundles, indicating successful insemination and sperm storage. Scale bars, 20 µm.

**Fig. S2.**

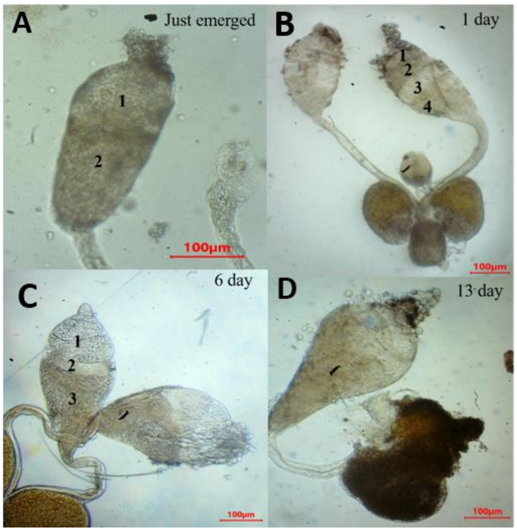

**Fig. S2. Age-dependent changes in testicular morphology of male *An. culicifacies***

(A-D) Representative bright-field microscopic images of testes dissected from <24hr, 1-day-old, 6-day-old, and 13-day-old male mosquitoes. Progressive reduction in the number of spermatocysts was observed with increasing age, indicating advancement of spermatogenesis and sperm maturation. Images were acquired using a light microscope at 10× magnification. Scale bars are indicated in the images.

**Fig. S3.**

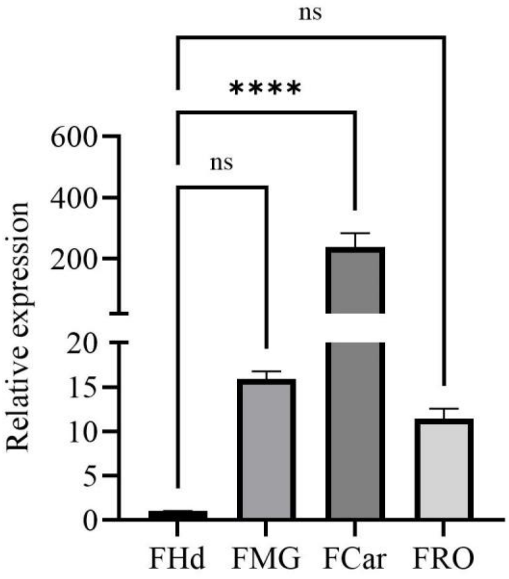


**Fig. S3. Tissue-specific expression profile of *AcLITAF6* in adult female mosquitoes**

Relative transcript abundance of *AcLITAF6* in the female head (FHd), carcass (FCar), midgut (FMG), and female reproductive organs (FRO) of 3–4-day-old naïve adult female determined by quantitative real-time PCR (qRT-PCR). Data represent mean (n = 25 mosquitoes per replicate, N = 3 biological replicates). The statistical significance relative to the female head (FHd), used as the control, is denoted as ns= not significant; P<0.0001 using one-way ANOVA with Dunnett's multiple comparisons test.

**Fig. S4.**

>ACUA022039-RA cds: protein_coding

ATGTCGAAAGATGGACCTCCACCGTACGGGTTCGTGCCACCGCCATCGGCACCA
CCAAGCTATGCGCAGGCTGTCGGTGGTGTACCACCGTCCAGCCCGTTCACACCGC
AGCAACCGGTACTGACCGGGGCACAGATTGTTACCACGGTCGTACCGATCGGAC
CACAGTCGACGCACATGGTATGTCCCAGCTGCCATGCGGAAGTTAACACCGAGA
CAACAACATCGCCGGGAATGATTGCTTACGTGTCCGGTTTCCTGATCGCACTGTT
TGGATGCTGGTTAGGATGCTGCCTAATACCGTGCTGCATTGACGAATGTATGGAT
ATCCATCATACCTGTCCACGCTGTCGAGCCTACCTGGGACGTCACAAGCGATAA

>ACUA022039-RA peptide: ACUA022039-PA pep:protein_coding

MSKDGPPPYGFVPPPSAPPSYAQAVGGVPPSSPFTPQQPVLTGAQIVTTVVPIGPQSTH
MVCPSCHAEVNTETTTSPGMIAYVSGFLIALFGCWLGCCLIPCCIDECMDIHHTCPRC
RAYLGRHKR

| Transcript ID | Transcript Length | Protein Length | Transcript Type | Genomic Length |
|---|---|---|---|---|
| ACUA022039-RA | 381 | 126 | mRNA | 1525 |

| Transcript ID | Isoelectric Point | Molecular Weight | Has SignalP | Has TMHMM | Protein Length |
|---|---|---|---|---|---|
| ACUA022039-RA | 7.21 | 13355 | no | yes | 126 |

**Fig. S4. Nucleotide sequence**

Coding DNA sequence (CDS) and predicted amino acid sequence of *AcLITAF6* (ACUA022039-RA) from *An. culicifacies*. The complete open reading frame and translated peptide sequence are shown to illustrate gene structure and predicted protein composition.

**Fig. S5.**

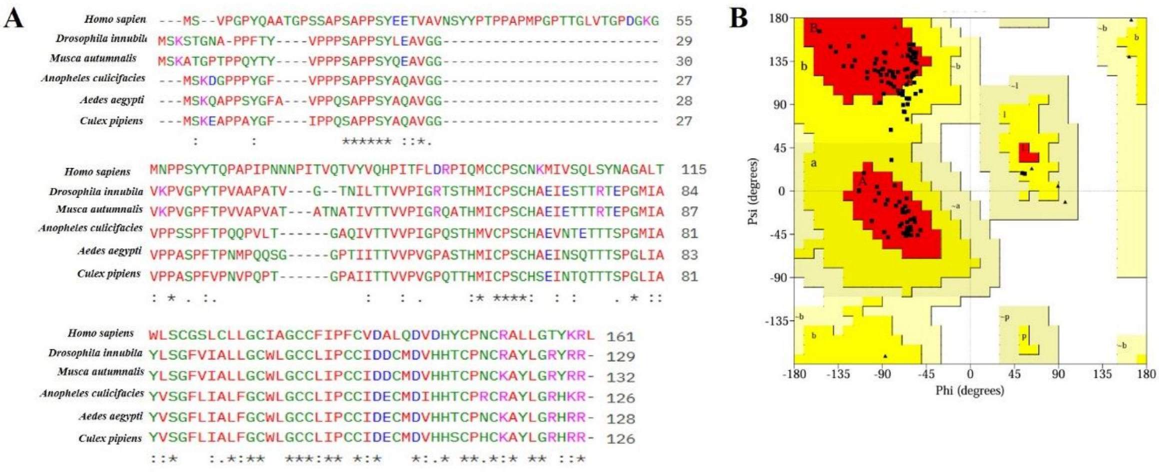


**Fig. S5. Multiple sequence alignment of *AcLITAF6* and Ramachandran plot**

(A) Multiple sequence alignment of *AcLITAF6* with selected LITAF homologues from representative insect species and humans, demonstrating conservation of key amino acid residues, particularly within the N-terminal region.

(B) Ramachandran plot of the predicted *AcLITAF6* protein structure showing stereochemical quality of the model, with 83.0% residues in favoured regions and 17.0% residues in allowed regions. Red, yellow, lemon, and white regions indicate favored, allowed, generously allowed, and disallowed conformations, respectively.

**Fig. S6.**

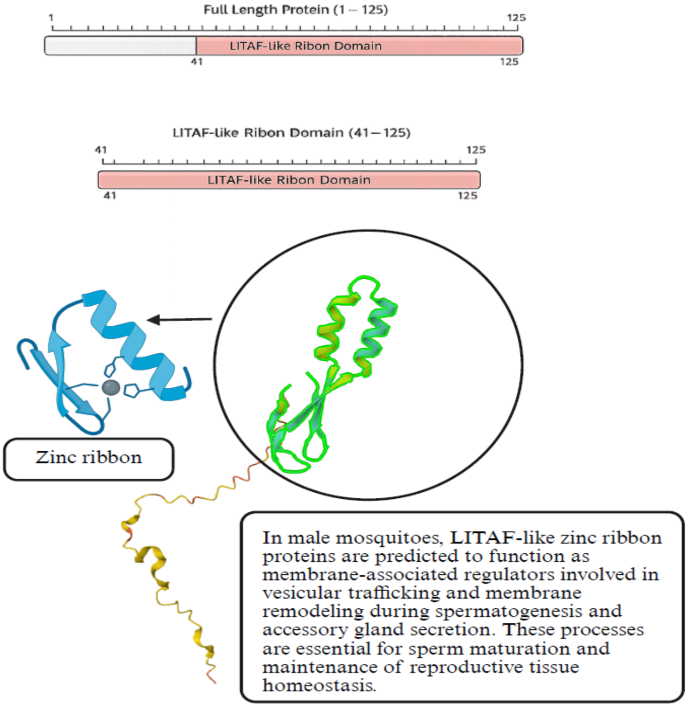


**Fig. S6. Structural organization and predicted functional role of the mosquito LITAF-like protein.**

Schematic representation of the mosquito LITAF-like protein showing the full-length amino acid sequence (1–125 aa) and the conserved LITAF-like zinc ribbon domain spanning residues 41–125.

**Fig. S7.**

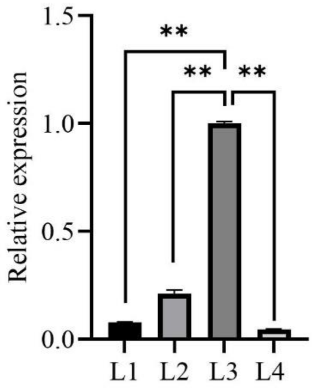

**Fig. S7. Relative expression analysis of *AcLITAF6* during the larval stages.**

Relative gene expression during larval development, showing higher expression of the transcript in L3 of the mosquito *An. culicifacies* (n=10, N=3). The statistical significance relative to the L3, used as the control, is denoted as P= 0.0029, P=0.0080, and P=0.0021 for L1, L2, and L4, respectively, using one-way ANOVA with Dunnett's multiple comparisons test.

**Fig. S8.**

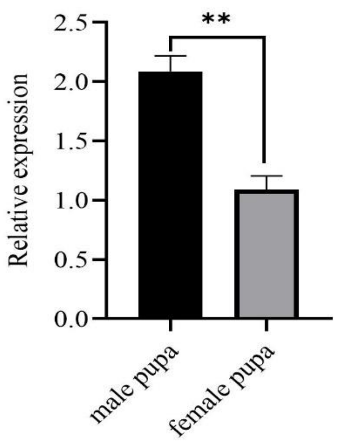


**Fig. S8. Relative expression analysis of *AcLITAF6* in male and female pupa**

Relative gene expression showing higher expression of the transcript in male pupa than female pupa of the mosquito *An. culicifacies* (n=10, N=3). The statistical significance relative to the female pupa, used as the control, is denoted as P=0.003 using an unpaired two-tailed Student's t-test.

**Fig. S9.**

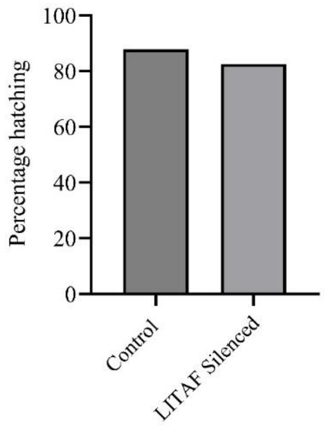


**Fig. S9. Effect of *AcLITAF6* silencing on egg hatching of mosquitoes.**

Percentage egg hatching was assessed in control and *AcLITAF6*-silenced females. Silencing of *AcLITAF6* resulted in a slight reduction in hatching rate compared with the control group; however, egg hatchability remained above 80% in both groups. Bars represent mean percentage hatching.

**Fig. S10.**

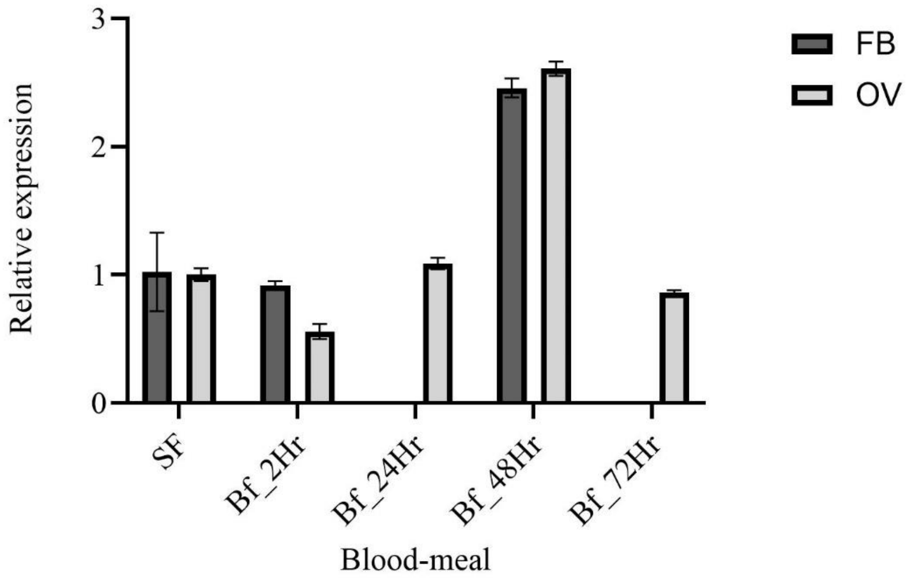


**Fig. S10. Relative expression analysis of *AcLITAF6* in fat body and ovary post blood meal** Relative expression of *AcLITAF6* in the fat body and ovaries of female mosquitoes following blood feeding. Transcript abundance increased progressively and reached maximal levels at 48 hr post-blood meal.

**Fig. S11.**

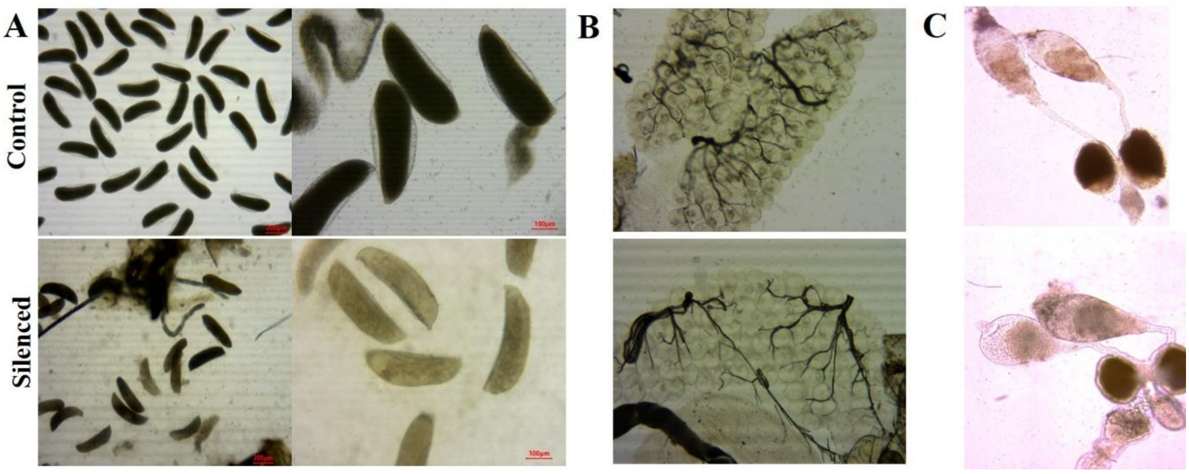


**Fig. S11. Microscopic images of reproductive tissues and egg laid**

(A) Microscopic images of eggs laid by female mosquitoes mated with control and *AcLITAF6*-silenced male mosquitoes at 4X and 10X magnification.

(B) Female reproductive organ (FRO) of control female mosquitoes mated with control and silenced male mosquitoes, showing nucleation in control. Images were taken at 10X magnification.

(C) Control and silenced the MRO of male mosquitoes, showing a compact sperm reserve in control as compared to the silenced MRO.

**Fig. S12.**

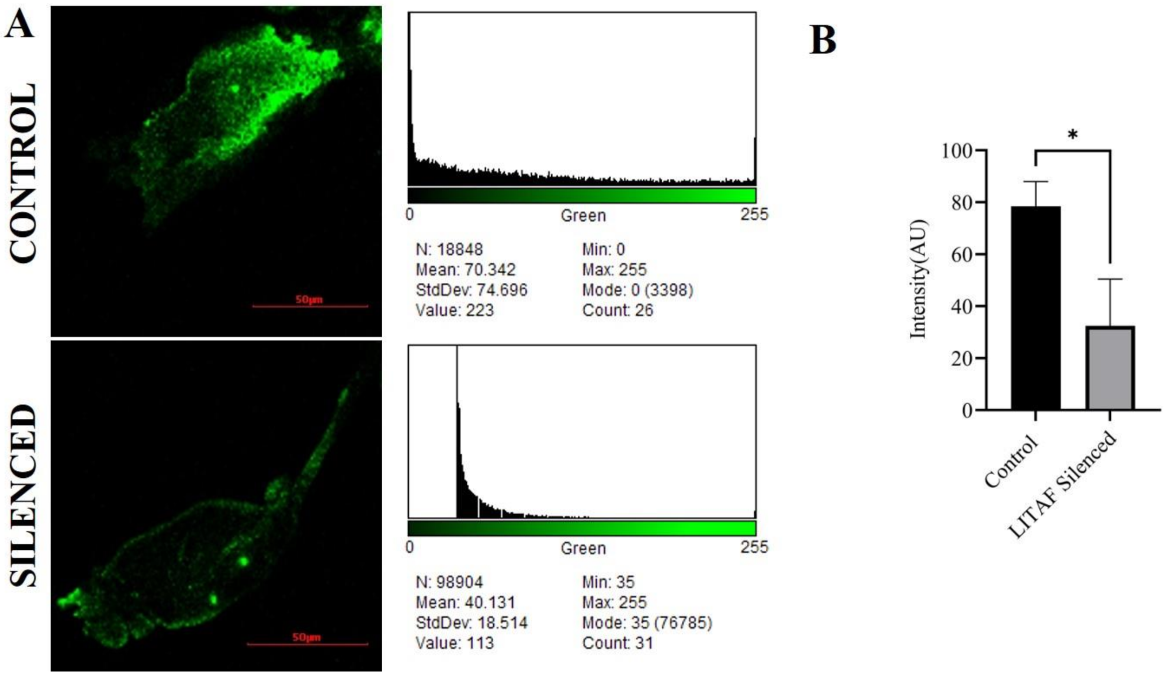


**Fig. S12. Quantification of apoptotic activity in male reproductive organs using TUNEL assay**

(A) Representative confocal micrographs and fluorescence intensity histograms showing apoptotic activity via TUNEL assay in control and *AcLITAF6*-silenced MROs.

(B) Fluorescence intensity corresponds to relative apoptotic activity. The statistical significance relative to the control, used as the reference, is denoted as P=0.017 using an unpaired two-tailed Student's t-test.

**Fig. S13.**

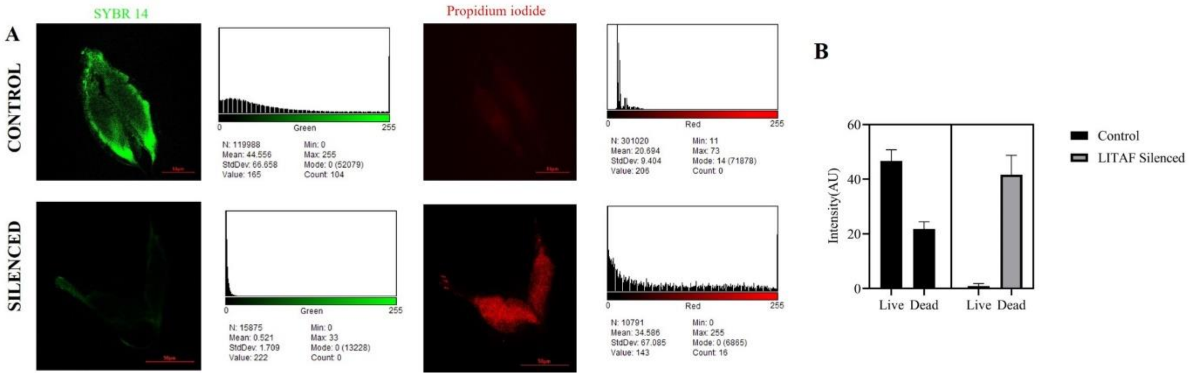


**Fig. S13. Quantification of live/dead in male reproductive organs using live/dead sperm viability assay**

(A) Representative confocal micrographs and fluorescence intensity histograms showing live/dead germ cells via live/dead sperm viability assay in control and *AcLITAF6*-silenced MROs.

(B) Fluorescence intensity corresponds to relative live and dead cells. The statistical significance relative to the control, used as the reference, is denoted as P=0.017 using an unpaired two-tailed Student's t-test.

**Fig. S14.**

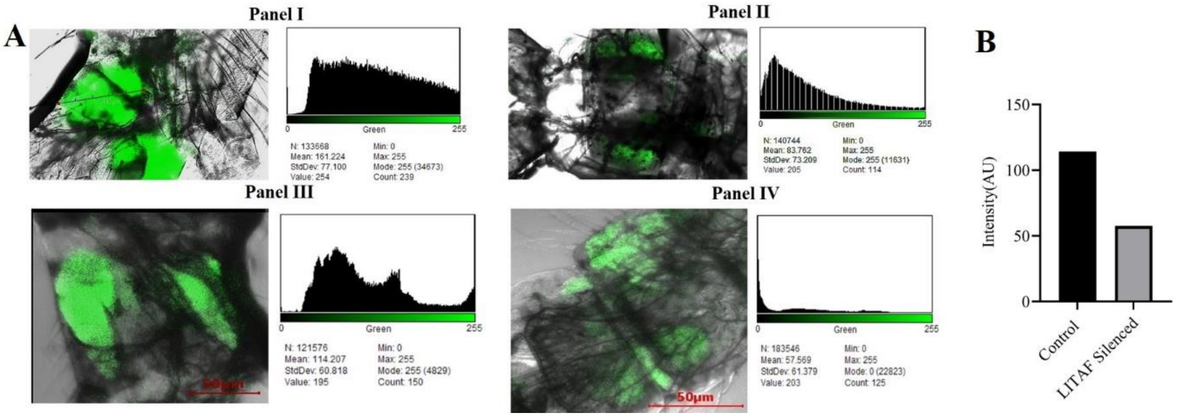


**Fig. S14. Quantification of phagocytic activity in male reproductive organs using pHrodo-labelled E. coli bioparticles**

Representative confocal micrographs and fluorescence intensity histograms showing phagocytic activity in male reproductive organs (MROs). Phagocytosis was assessed using pHrodo Red E. coli bioparticles, which emit fluorescence upon internalization into acidic phagosomes. Panels represent virgin MROs (Panel I), mated MROs (Panel II), control dsGFP-injected MROs (Panel III), and *AcLITAF6*-silenced MROs (Panel IV). Fluorescence intensity corresponds to relative phagocytic activity.

**Fig. S15.**

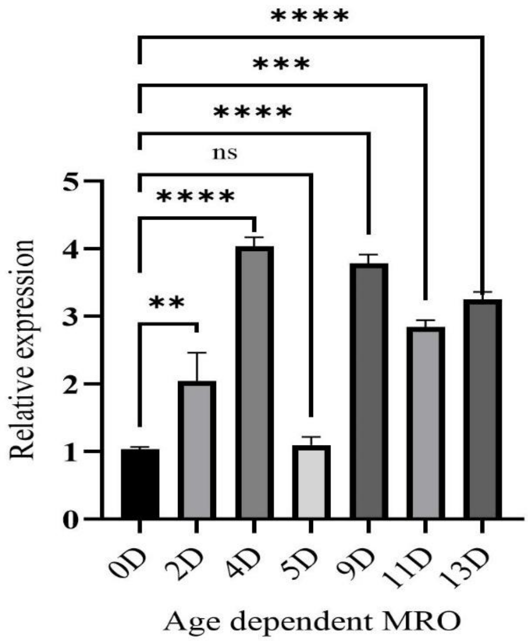


**Fig. S15. Age-dependent expression pattern of *AcLITAF6* in male reproductive organs**

Relative expression of *AcLITAF6* in MROs collected from <24hr, 1-day-old, 4-day-old, 9-day-old, and 13-day-old virgin male mosquitoes. Transcript levels were quantified by qRT-PCR, normalized with AcRSP7, and calculated using the $2-\Delta\Delta Ct$ method. Data represent mean (n = 25 mosquitoes per replicate, N = 3 biological replicates). Statistical significance was determined by one-way ANOVA followed by Dunnett's multiple-comparisons test using <24hr as the reference group. ns = not significant, ** $p < 0.01$, *** $p < 0.001$, **** $p < 0.0001$.

**Video S1 Microscopic video of sperm in the control and silenced testes of male mosquitoes.** Video showing less movement of sperm in silenced testes than in control testes.

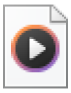

CONTROL.wmv

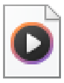

SILENCED.mp4
